# Conserved Molecular Responses to Arsenite Exposure in *Drosophila melanogaster*

**DOI:** 10.64898/2026.02.08.701855

**Authors:** Shannon R. Smoot, Jay Tourigny, Jessica M. Holsopple-Bowen, Alexander J. Fitt, Yadira Pena-Garcia, Matt R. Lowe, Darcy A. Rose, Noelle C. Zolman, Grace H. Thrasher, Satyavsh S. Arya, Hunter B. Vires, Emma C. Adams, John K. Colbourne, Joseph R. Shaw, Brian Oliver, Travis Nemkov, Angelo D’Alessandro, Thomas C. Kaufman, Jason M. Tennessen

**Author notes:** Equal Contributions (Shannon R. Smoot, Jay Tourigny).

## Abstract

Arsenic exposure is a pervasive global health threat strongly associated with increased risk of morbidities such as diabetes, cardiovascular disease, and cancer. Despite extensive studies describing the dangers of arsenic exposure, the molecular initiating events that link arsenic to chronic disease onset and progression remain poorly defined. To address this knowledge gap, we combined time-resolved transcriptomic and metabolomic profiling of adult *Drosophila melanogaster* exposed to sodium (meta) arsenite (NaAsO_2_). We uncovered coordinated, dose-dependent shifts in gene expression and metabolite abundance that activate canonical detoxification pathways and mirror arsenic-associated disease signatures in humans. Notably, flies rapidly upregulated heatshock and xenobiotic response gene networks, followed by biomarkers characteristic of diabetic states (elevated glucose, lactate, and methylglyoxal, for example). These findings reveal conserved molecular pathways that couple arsenic exposure to metabolic dysfunction and establish *Drosophila* as a powerful whole-organism model for identifying early biomarkers and mechanistic drivers of arsenic-induced disease phenotypes.

## INTRODUCTION

Over 200 million people worldwide are exposed to arsenic through multiple environmental and dietary sources, posing a substantial global public-health concern (Organization, 2022, Podgorski and Berg, 2020). Arsenic contamination is especially severe in the heavily populated regions of Bangladesh and West Bengal, where naturally high concentrations occur in soil and groundwater (Chowdhury et al., 2000, Mazumder et al., 2010, Ivy et al., 2023). Similarly, arsenic contamination is a major health hazard for the western and midwestern regions of the United States, as well as parts of northern New England, where groundwater in millions of domestic wells exceeds the World Health Organization (WHO) drinking water guideline of 10 μg/L (Organization, 2022, Ayotte et al., 2017).

In addition to drinking water, the global food-supply system distributes arsenic exposure far beyond these regions, adding to overall intake and increasing cumulative exposure risk. Rice (*Oryza sativa* L.) is particularly vulnerable to arsenic contamination, often containing concentrations up to ten times higher than other domesticated crops such as wheat or corn (Su et al., 2010, Upadhyay et al., 2019). Consequently, populations that rely on rice as a staple food may consume significant quantities of arsenic, as do infants and young children fed rice-based baby formula and baby foods (Signes-Pastor et al., 2016, Parker et al., 2022, Wang et al., 2025). The risk of arsenic exposure similarly extends to populations that consume large amounts of seafood, as some marine species, such as shellfish, bioaccumulate arsenic at levels that can substantially exceed those found in most terrestrial foods, in some cases approaching or exceeding levels associated with adverse health outcomes (Taylor et al., 2017).

Importantly, these dietary exposures add to arsenic intake from drinking water and other sources, increasing cumulative exposure risk. Together, these diverse and overlapping exposure routes complicate efforts to mitigate arsenic risk at its source and underscore the need to understand how arsenic perturbs conserved biological pathways. A deeper understanding of the molecular responses to arsenic exposure is therefore critical for identifying biomarkers and molecular initiating events that precede the development of disease phenotypes associated with arsenic exposure.

Arsenic exposure results in pleiotropic health effects, as both acute and chronic exposures affect multiple organ systems and increase the risk of serious disease. The mechanisms underlying acute arsenic toxicity are relatively well defined and include disruption of mitochondrial oxidative phosphorylation, leading to decreased ATP production, increased reactive oxygen species, and activation of cell death pathways (Hughes, 2002). In contrast, chronic arsenic exposure, which more commonly occurs through ingestion of contaminated water, food, or air, is associated with a broad spectrum of long-term health outcomes, including disorders of the skin, lungs, liver, gastrointestinal tract, cardiovascular system, kidneys, and nervous system (Jomova et al., 2011, Singh et al., 2011, Tchounwou et al., 2019). Moreover, prolonged exposure significantly raises the risk of developing diabetes mellitus as well as cancers of the skin, liver, lungs, urinary bladder, and kidney (Lai et al., 1994, Ratnaike, 2003, Sharma and Sohn, 2009, Hughes, 2002, Singh et al., 2015). Thus, while the cellular consequences of acute arsenic toxicity are well characterized, a deeper understanding of the molecular initiating events and adaptive responses that link arsenic exposure to chronic disease onset and progression remains a critical unmet need. Given the widespread nature of arsenic exposure and its diverse health impacts, new genetic models are essential for rapidly dissecting the molecular mechanisms underlying systemic arsenic toxicity in a cost-efficient and ethical manner.

The fruit fly *Drosophila melanogaster* is an important genetic model to investigate the biological effects of arsenic exposure on animals, including humans, with previous studies examining how arsenic disrupts development, neurobiology, metabolism, and fertility (Goldstein and Babich, 1989, Ramos-Morales and Rodriguez-Arnaiz, 1995, Adebambo et al., 2024, Anushree et al., 2023, Anushree et al., 2022, Rizki et al., 2002, Rizki et al., 2006, Bahadorani and Hilliker, 2009). In addition, transcriptomic and population genetic studies in the fly have identified phylogenetically conserved mechanisms that influence arsenic sensitivity, underscoring the power of using *Drosophila* to identify and characterize the molecular mechanisms that control toxicity responses (Muñiz Ortiz et al., 2009, Muñiz Ortiz et al., 2011, Rand, 2010, Rand et al., 2023, Sykiotis and Bohmann, 2008, Vincent and Tanguay, 1982, Chaturvedi et al., 2025). However, much of this work has focused on individual genes or single exposure time points, leaving the temporal dynamics of arsenic-induced changes in gene expression and metabolite levels—particularly during the earliest stages of exposure— poorly defined. Because these immediate molecular responses likely represent initiating events that precede overt pathology, resolving how transcriptional and metabolic programs evolve over time is essential for understanding how arsenic exposure progresses toward disease states. To address this gap, we applied a multi-omic approach to track how the molecular response to arsenic evolves over time. Using semi-targeted UHPLC-MS-based metabolomics and RNA-seq, we comprehensively characterized transcriptomic and metabolic changes in *Drosophila* following NaAsO_2_ exposure across a 48-hour time course.

This integrated analysis revealed dose- and time-dependent shifts in gene expression and metabolite abundance that mirror patterns observed in mammalian systems and align with molecular signatures previously associated with arsenic-related human disease, indicating that these responses are evolutionarily conserved. Notably, we observed enrichment of differentially expressed genes linked to diabetes mellitus across sexes and exposure levels, even during this short-term exposure paradigm.

Consistent with these transcriptional changes, metabolomic profiling uncovered disruptions in carbohydrate metabolism and purine degradation that resemble metabolic features commonly associated with diabetic states. Together, these findings highlight conserved molecular responses that emerge rapidly following arsenite exposure and underscore the value of *Drosophila* as a model for identifying early biomarkers and molecular initiating events associated with chronic arsenic-related disease, rather than overt disease phenotypes themselves.

## METHODS

### *Drosophila* Genetics and Husbandry

Fly stocks were maintained on Bloomington *Drosophila* Stock Center (BDSC) media (https://bdsc.indiana.edu/information/recipes/bloomfood.html) at 25ºC. Unless noted, all studies described herein used Oregon-R (RRID:BDSC_25211). Flybase (https://flybase.org/) was utilized as a resource throughout these studies (Öztürk-Çolak et al., 2024).

### Dose-response curves

A stock solution of sodium (meta)arsenite (NaAsO_2_, Sigma-Aldrich, St. Louis, MO) was prepared by dissolving an accurately massed portion of NaAsO_2_ into a volumetrically measured amount Milli-Q water. The solution was stirred for 15 minutes until complete dissolution and titrated to neutral pH (∼7.5) with concentrated HCl. Dose-response curves were performed as previously described (Holsopple et al., 2023).

Briefly, bottle cultures containing Oregon-R male and female flies were allowed to eclose for two days at 25ºC, and the resulting adults were transferred to bottles containing fresh BDSC food. Following a two-day incubation at 25ºC, adult male and female flies were sorted by sex and aged in vials of BDSC food for an additional 48 hours at 25ºC to allow for recovery from the CO_2_ anesthesia used for sorting. Flies within individual vials were then transferred without anesthesia to starvation vials (Whatman filter paper No. 1 soaked in sterile milli-Q water) for 16 hours at 25°C. Following the starvation period, flies were transferred without anesthesia to exposure vials containing liquid food [4% sucrose (m/v), 1.5% yeast extract (m/v)] and the noted concentration of NaAsO_2_. The number of dead female or male flies per vial was counted in each vial at 24 and 48 hours. A total of six replicate vials were scored for each concentration. Dose-response curves were calculated using the frequentist dichotomous Hill model in Benchmark Dose Software (BMDS), published by the U.S. Environmental Protection Agency (Agency, 2023). The dose response curves between males and females were analyzed with GraphPad Prism 10.5.0 using the Dose-Response Special ECanything equation.

### Sample Collection

NaAsO_2_-exposed flies and unexposed controls were prepared for LC-MS and RNA-seq using the same protocol described for the dose-response curves (Holsopple et al., 2023). Briefly, following the overnight starvation, vials of 20 male or female flies were transferred into a vial containing liquid media supplemented with 0 mM, 0.25 mM, or 1.0 mM NaAsO_2_. Samples were then collected at 1, 2, 4, 8, 24, and 48 hrs by individually transferring the flies from the exposure vials to empty fly culture vials, which were then immediately placed on dry ice. Once immobilized, the number of flies in individual vials was counted and subsequently transferred to a tared 2 mL screw cap tube containing 1.4 mm ceramic beads (Fisher Brand; 15-340-153). The cap was screwed onto the tube, the sample mass measured using a Radwag AS 82/220.X2 Plus analytical balance, and the closed tube immediately placed in liquid nitrogen. Frozen tubes were stored in an -80ºC freezer.

### RNA preparation

Briefly, bead tubes containing either control or exposed flies were transferred from the –80ºC freezer into a tube rack surrounded by dry ice. 800 μL of Trizol reagent (Invitrogen; 15596018) was added to each tube and the sample was homogenized in a 4ºC room for 30 secs at 6.45 m/sec using an Omni Beadruptor 24. Homogenized samples were removed from the instrument and immediately placed on ice for 2 min, then allowed to incubate for 5 min at room temperature. After incubation, 160 μL of chloroform was added to each tube and vortexed for 15 seconds. Samples were then incubated at room temperature for 15 min and subsequently centrifuged at 12,000 x g for 15 min at 4°C. Following centrifugation, 300 μL of the upper aqueous phase was transferred to an RNase-free 1.5 mL microfuge tube containing 400 μL of isopropanol, mixed by inverting, and centrifuged at 12,000 x g for 15 min at 4°C. The isopropanol was removed from each sample using a 200 μL pipette while being careful not to disturb the RNA pellet. 1 mL of 75% ethanol was added to each tube and samples were centrifuged at 12,000 x g for 15 min at 4°C. The ethanol was removed from each sample using a 200 μL pipette while being careful not to disturb the white RNA pellet.

The samples were allowed to briefly air dry to completely remove the remaining ethanol. Samples were resuspended in 50 μL of RNAase free water and stored in a -80°C freezer. Resuspended RNA samples were subsequently cleaned using a Qiagen RNeasy Mini Kit per manufacturer’s instructions (Qiagen; 74104).

### RNA sequencing

Libraries of these 114 samples, 57 of each sex, were generated with the Illumina TruSeq Stranded mRNA HT protocol and were loaded evenly on the NextSeq 2000 P3 platform and run for 100 cycles, with a target of ≈8.8 million 2×50 paired-end reads per sample. Any samples with < 9 million reads were topped up in a further run, with additional reads concatenated to their respective samples. Raw reads and processed counts are available on NCBI GEO at accession GSE241663. All library preparation and sequencing were performed at the IU Center for Genomics and Bioinformatics (CGB).

### Transcriptomic Analysis

RNA-seq read quality was assessed with FastQC (Andrews, 2010) and MultiQC (Ewels et al., 2016); raw reads were not trimmed or filtered. Reads were quasi-aligned and quantified using Salmon v1.10.2 (Patro et al., 2017), with the Salmon index built from the *D. melanogaster* BDGP6.32 transcriptome and reference assembly (for decoy sequences) retrieved through Ensembl (Yates et al., 2021).

Differential expression analysis was performed with tximport (Soneson et al., 2015) and DESeq2 v1.30.1 (Love et al., 2014) running in RStudio v2023.07.999 on R v4.3.1. For count tables, including raw counts, normalized counts, and variance-stabilized log counts, significance of differentially expressed genes was assessed with the likelihood-ratio test (LRT), adjusted *p* ≤ 0.05. For pairwise comparisons within sex at each time point and arsenite concentration relative to its time-matched 0 mM control, the Wald test with a concatenated contrast of “[sex]_[concentration]_[time]” vs. “[sex]_0mM_[time]” was used to identify genes with adjusted *p* ≤ 0.05 and an absolute log fold change (abs(LFC)) ≥ 1. All computation was performed on Indiana University’s High Performance Computing clusters using system modules and user conda environments.

### Gene Set Enrichment Analysis

The list of genes that were significantly altered in 0.25 mM and 1.0 mM NaAsO2-exposed flies compared to 0 mM controls was analyzed using PAthway, Network and Gene-set Enrichment Analysis (PANGEA) (Hu et al., 2023) using the “Search Multiple Gene Lists” function. PANGEA analyses were conducted as follows: (i) the time course analysis used SLIM GO BP, FlyBase Signaling Pathway (Experimental Evidence), KEGG Pathway D.mel, and REACTOME Pathway; (ii) the disease enrichment analysis used FlyBase phenotypes for classical alleles and Disease Annotation AGR; (iii) tissue expression analysis used Preferred tissue (modEncode RNA_seq) and Expression annotation AGR. All results are presented as heatmaps with GO categories for the time point of interest rank-ordered according to adjusted *p*-value. PANGEA was also used to analyze the 100 genes that are differentially expressed in all four timecourses (male & female, 0.25mM & 1mM arsenite) using the parameters described above. The node graph was generated in PANGEA based on the top 20 most significantly enriched gene sets. Gene sets with the lowest amount of redundancy were selected to generate the graph.

### LC-MS Sample Preparation

For metabolomic analysis, frozen samples were transferred from a -80°C freezer to a -20°C-cooled enzyme carrier caddy. 800 μL of prechilled (-20°C) 90% methanol containing 2 μg/mL succinic-d4 acid was added to each sample tube using a positive displacement pipette. The samples were homogenized using an Omni Beadruptor 24 (Omni International) at 4°C for 30 seconds at 6.45 m/sec. Once homogenized, the samples were incubated at -20°C for 2 hours. After incubation, the samples were centrifuged at 12,000 x g for 5 min at 4°C. The tubes were carefully removed from the centrifuge and 600 μL of the upper supernatant was transferred to a new labelled 1.5 mL microcentrifuge tube on dry ice. Each tube containing the newly extracted supernatant was centrifuged 12,000 x g for 5 min at 4°C. 200 μL of supernatant from each sample was transferred to a designated well in a 96 well plate on dry ice. Once the plate was fully loaded with sample supernatants, the plate was dried under a nitrogen evaporator. Once fully dried, the plate was covered with an aluminum seal and stored at -80°C. Remaining supernatant was transferred to new labelled 1.5 mL microcentrifuge tubes and dried down and stored at -80°C.

### Ultra High-pressure Liquid Chromatography - Mass Spectrometry (UHPLC-MS)-based Metabolomics

UHPLC-MS metabolomics analyses were performed at the University of Colorado Anschutz Medical Campus, as previously described (Nemkov et al., 2019). Briefly, the analytical platform employs a Vanquish UHPLC system (Thermo Fisher Scientific, San Jose, CA, USA) coupled online to a Q Exactive mass spectrometer (Thermo Fisher Scientific, San Jose, CA, USA). The (semi)polar extracts were resolved over a Kinetex C18 column, 2.1 x 150 mm, 1.7 μm particle size (Phenomenex, Torrance, CA, USA) equipped with a guard column (SecurityGuard™ Ultracartridge – UHPLC C18 for 2.1.0 mM ID Columns – AJO-8782 – Phenomenex, Torrance, CA, USA) using an aqueous phase (A) of water and 0.1% formic acid and a mobile phase (B) of acetonitrile and 0.1% formic acid for positive ion polarity mode, and an aqueous phase (A) of water:acetonitrile (95:5) with 1.0 mM ammonium acetate and a mobile phase (B) of acetonitrile:water (95:5) with 1.0 mM ammonium acetate for negative ion polarity mode. The Q Exactive mass spectrometer (Thermo Fisher Scientific, San Jose, CA, USA) was operated independently in positive or negative ion mode, scanning in Full MS mode (2 μscans) from 60 to 900 m/z at 70,000 resolution, with 4 kV spray voltage, 45 sheath gas, 15 auxiliary gas. Calibration was performed prior to analysis using the Pierce™ Positive and Negative Ion Calibration Solutions (Thermo Fisher Scientific).

### Statistical Analysis of Metabolomics Data

All metabolomics datasets were analyzed using Metaboanalyst 5.0 (Pang et al., 2021), with data normalized to sample mass and preprocessed using log normalization and Pareto scaling. Data generated by GC-MS analysis of individual compounds was analyzed using GraphPad Prism 10.

### Data Availability

All strains and reagents are available upon request. Processed RNA-seq data is presented in Tables S2-S9 and raw data are available in NCBI Gene Expression Omnibus (GEO; GSE241663). All metabolomics data described herein are included in Tables S10 and S11.

## RESULTS

### NaAsO_2_ exposure induces conserved transcriptional responses in adult *Drosophila*

To investigate dynamic molecular responses activated in *Drosophila* adults upon NaAsO_2_ exposure, we used a combined transcriptomic and metabolomic approach to investigate time- and dose-dependent changes in gene expression and metabolite abundance. Previous studies using 48 hr NaAsO_2_ exposures (Holsopple et al., 2023, Adebambo et al., 2024, Hayot et al., 2025), as well as our own analysis (Figure S1, Table S1), established LD_10_ values for Oregon-R males and females ranging between 0.3 mM and 0.5 mM, and LD_50_ values ranging from 0.65 mM to 1.0 mM. Based on these benchmarks, we exposed Oregon-R adult male and female flies to sublethal (0.25 mM; <LD_10_) and lethal (1.0 mM; >LD_50_) NaAsO_2_ concentrations, aiming to capture both adaptive responses that help maintain homeostasis and pathological responses. Samples were collected at 1, 2, 4, 8, 24, and 48 hours of exposure for RNA-seq analysis (Table S2-9), and 2, 4, 8 24, and 48 hrs for semi-targeted UHPLC-MS-based metabolomic analysis (Table S10-11). For each time point, changes in gene expression and metabolite abundance were compared to sex-matched, time-matched unexposed controls.

Our initial analysis of both the transcriptomic and metabolomic data revealed dose-, time-, and sex-dependent differences in response to NaAsO_2_ (Figures 1, 2, and S2; Table S12 and S13). We noted immediately that the sex-specific differences are extensive, with less than half of the DEGs shared between males and females at any given time point (Figure S2). These sex-specific changes in gene expression and metabolite levels will be addressed in detail elsewhere (Tourigny *et al*., In preparation). In this study, we focused on the dose- and time-dependent molecular responses shared between NaAsO_2_-exposed males and females.

**Figure 1.**
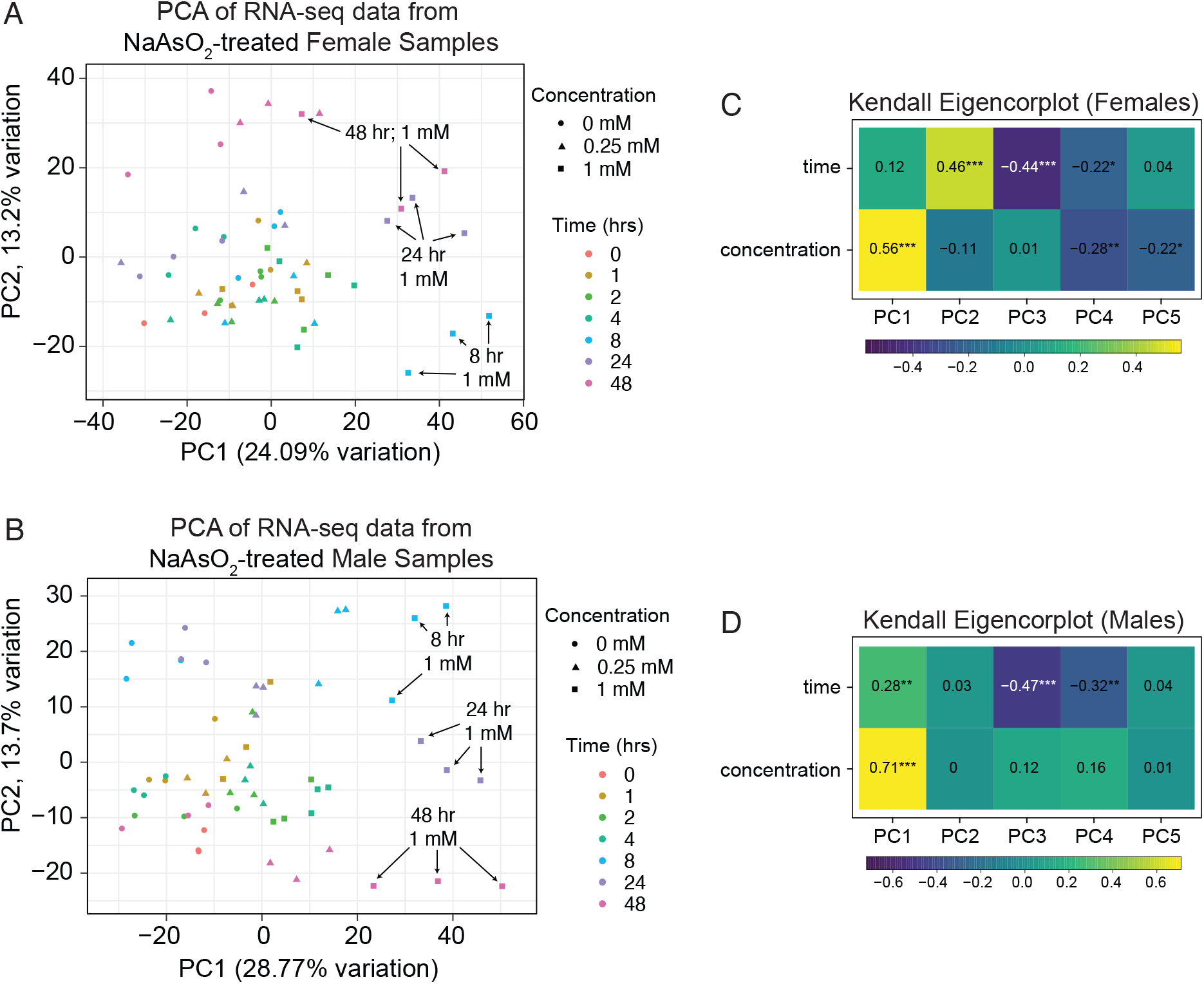
Principal Component Analysis (PCA) of RNA-seq data from adult *Drosophila* exposed to NaAsO_2_ across a 48-hour time course. PCA was used to analyze RNA-seq data from adult (A) male and (B) female flies exposed to control diet, 0.25 mM NaAsO_2_, and 1.0 mM NaAsO_2_. In both sexes (A,B), samples exposed to 1.0 mM mM NaAsO_2_ for 8, 24, and 48 hrs separated clustered in a similar position along the PC1 axis. RNA-seq data are available in Table S2, S4, S6, and S8.

**Figure 2.**
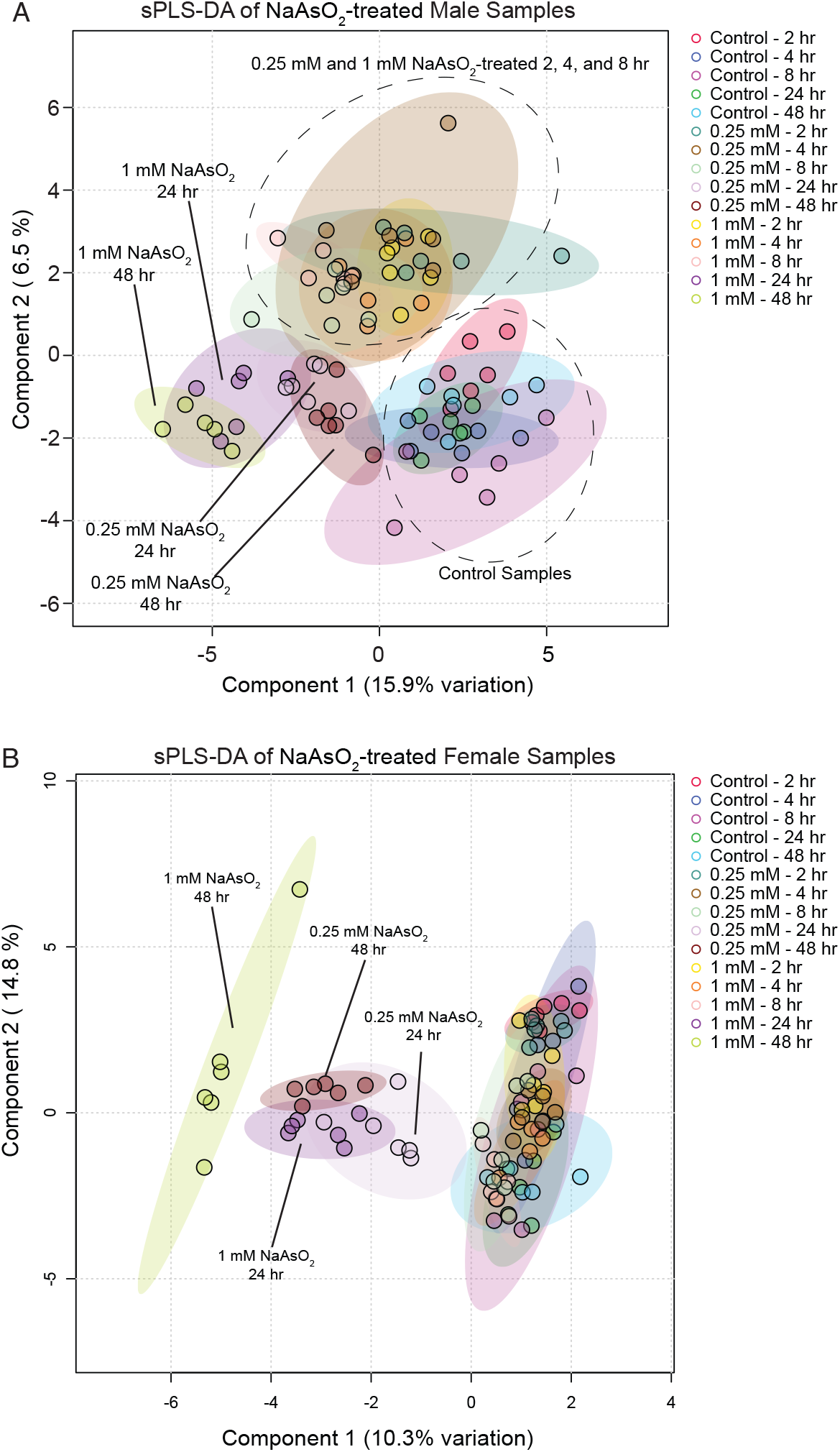
Partial Least Squares Discriminant Analysis (PLS-DA) of metabolomics data from adult *Drosophila* exposed to NaAsO_2_ across a 48-hour time course. Metaboanalyst 6.0 was used to conduct a PLS-DA of metabolomics data from adult (A) female and (B) male flies exposed to control diet, 0.25 mM NaAsO_2_, and 1.0 mM NaAsO_2_. In (A), the control samples and time points 2, 4, and 8 hr group compose two distinct clusters and are encircled by a dashed line. In (B), the 24 and 48 hr time points from both the low- and high-dose exposure cluster separately from all other sample sets. (C,D) Heatmap of Kendall’s Tau correlations of the experimental variables with principal components 1 through 5 for (C) females and (D) males. Asterisks denote increasing levels of statistical significance. Analyzed data are available in Table S10 and S11.

To compare the kinetics of transcriptional and metabolic responses to arsenite, we performed Principal Component Analysis (PCA) on the RNA-seq data and PLS-DA on the metabolomics. In the RNA-seq data, samples separated primarily along PC1 (Figure 1A–B), which was largely driven by arsenite concentration, with additional contributions from time (Figures 1C-D). By 8 hours of exposure, transcriptional profiles, particularly at 1.0 mM, had already diverged from earlier time points, with samples from 8, 24, and 48 hours clustering together along PC1. In contrast, metabolomic profiles showed little separation at 8 hours, with distinct clustering emerging only at 24 and 48 hours (Figure 2A-B). Together, these analyses indicate that arsenite induces rapid, dose-dependent transcriptional changes that precede large-scale metabolic remodeling.

Because arsenite concentration emerged as the dominant driver of transcriptional variation, we focused first on the RNA-seq data to identify dose-dependent gene expression programs that precede metabolic changes. Using PANGEA (Hu et al., 2023), we performed gene set enrichment analysis across all time points and exposure conditions (Table S14). Examination of the top 10 most significantly enriched gene sets at each time point revealed dynamic gene expression programs associated with metabolic, xenobiotic, and stress responses (Figures 3, 4, S3, S4 and Table S14). Strikingly, all datasets showed early enrichment for targets of the Heat Shock Transcription Factor (Hsf) (Clos et al., 1990, Zimarino et al., 1990, Wu et al., 1987), with “HSF-1-dependent transactivation” (R-DME-3371571) and “Regulation of the HSF1-mediated heat shock response” (R-DME-3371453) ranking among the most significantly enriched gene sets at either the 1 hr or 2 hr time point in all four datasets. Consistent with these findings, NaAsO_2_ elicits induction of Heat-shock protein transcripts following as little as 1 hr of exposure (Figure 5A-B, and Table S14), highlighting the Hsf gene regulatory network as one of the earliest responders to NaAsO_2_-exposure. We hypothesize that this response, although strongly induced by arsenite in our system, is unlikely to be arsenic-specific and instead represents a conserved stress response shared across diverse toxic exposures.

**Figure 3.**
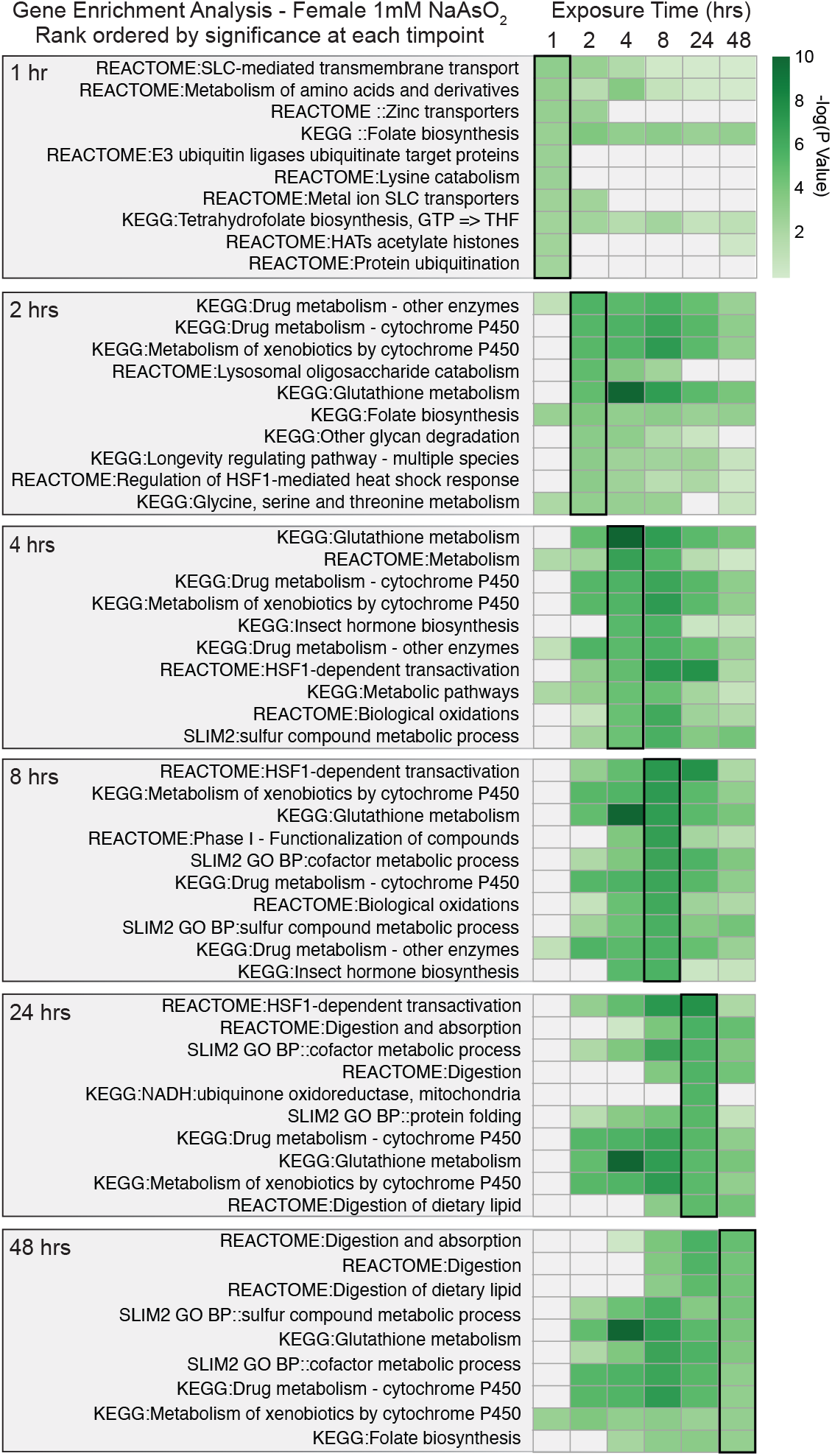
Gene Set Enrichment Analysis of DEGs in female *Drosophila melanogaster* exposed to 1.0 mM NaAsO_2_ across a 48-hour time course. PANGEA was used to identify significantly overrepresented gene sets among differentially expressed genes (DEGs) at each exposure time point (see Methods and Table S14). The top 10 enriched gene sets for each time point are presented with heatmaps of their rank-ordered adjusted *p*-value at the corresponding time (outlined heat map column), along with their significance (green gradient) or lack thereof (grey) at the other time points.

**Figure 4.**
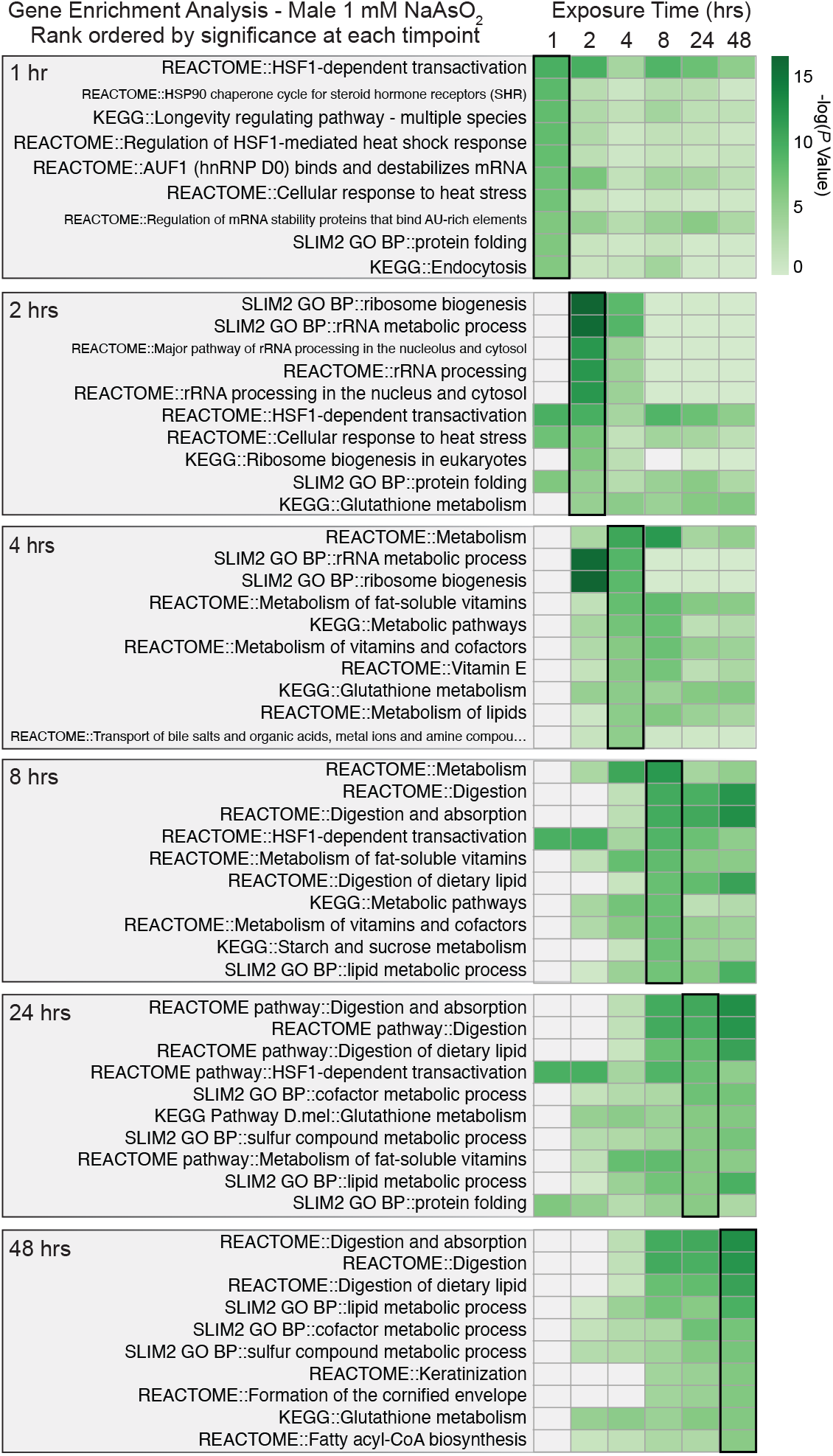
Gene Set Enrichment Analysis of DEGs in male *Drosophila melanogaster* exposed to 1.0 mM NaAsO_2_ across a 48-hour time course. As in Figure 3, PANGEA was used to identify significantly overrepresented gene sets among differentially expressed genes (DEGs) at each exposure time point (See Methods and Table S14). The top 10 enriched gene sets for each time point are presented with heatmaps of their rank-ordered adjusted *p*-value at the corresponding time (outlined heat map column), along with their significance (green) or lack thereof (grey) at the other time points.

**Figure 5.**
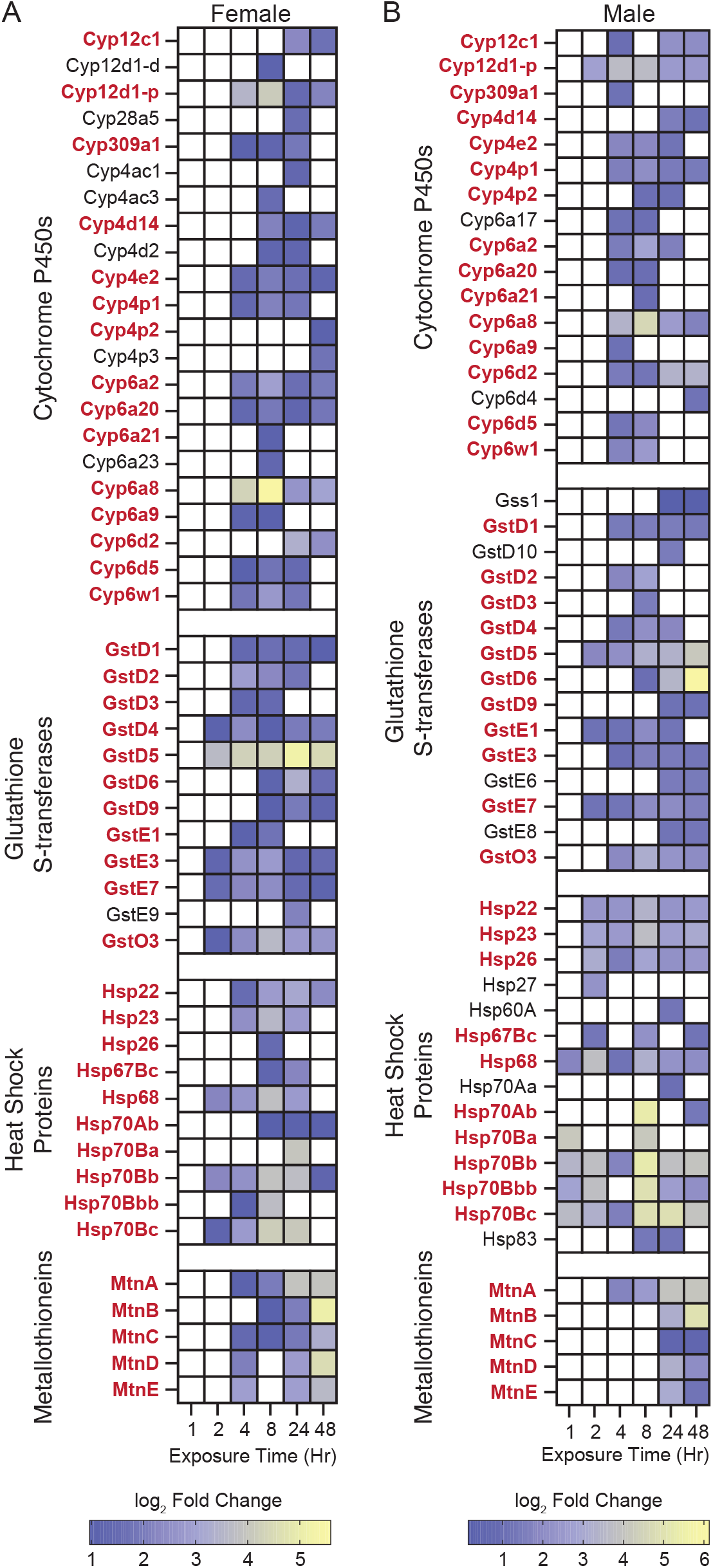
Up-regulated gene families across a 48 hr time course in male and female *Drosophila melanogaster* exposed to 1.0 mM NaAsO_2_. (A, B) Heatmaps displaying log_2_ fold changes of pairwise comparisons (time-matched treatment vs. control) for significantly upregulated (adj *p* ≤ 0.05) genes belonging to four canonical detoxification and stress-response families: cytochrome P450s (FBgg0001222), glutathione S-transferases (FBgg0000077), heat shock proteins (FBgg0000501), and metallothioneins (FBgg0000197). Gene expression changes are presented separately for (A) females and (B) males across 6 exposure time points (1, 2, 4, 8, 24, and 48 hours). Gene names in **bold** red exhibit significant up-regulation in both sexes. Non-significant time points are left blank. The data reveal distinct temporal dynamics and sex-specific patterns of gene induction among the canonical detoxification response pathways. Notably, males exhibit earlier activation of heat shock proteins, with females showing more rapid upregulation of metallothioneins. These findings highlight the coordinated and sex-dependent transcriptional activation of detoxification pathways in response to arsenite exposure.

Beyond the early heat shock response, several additional gene sets associated with metabolic and stress pathways were significantly enriched during the first 8 hours of exposure (Figures 3, 4, S3, S4, and Table S14). These gene sets are primarily composed of cytochrome P450s (path:map00982; path:map00980), glutathione S-transferases (Gst; path:map00480), and metallothioneins (Mtn; GO:0042221)—all key components of arsenic and metal detoxification networks in flies and humans (Yiwen et al., 2022, Muñiz Ortiz et al., 2009, Thomas, 2009). A closer examination of the significant pairwise comparisons revealed that these three enzyme families are coordinately upregulated throughout the 48 hrs of exposure (Figure 5A-B and S5A-B), although the exact sets of genes activated at these time points varied by dose and sex.

At later time points, gene sets linked with digestion and lipid metabolism became prominent (Figures 3, 4, S3, S4, and Table S14), suggesting that intestinal gene expression networks are especially sensitive to arsenite with continuous exposure. Consistent with this, tissue-specific enrichment analysis identified the digestive system as the most strongly associated tissue across all conditions (Figure 6A and Table S15).

**Figure 6.**
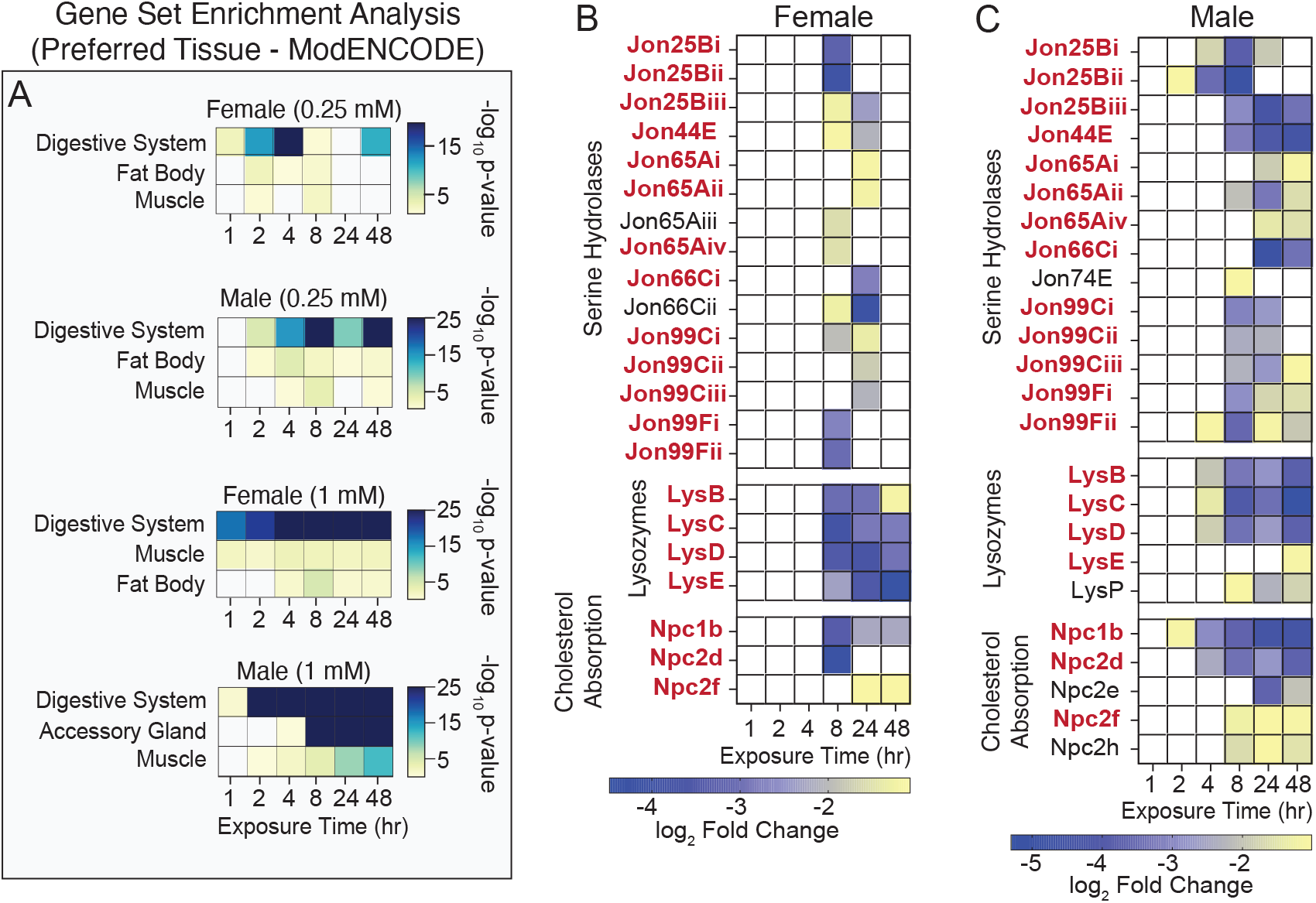
Identification of tissue-associated transcriptional responses to NaAsO_2_ across a 48-hour time course. (A) PANGEA was used to identify enrichment for tissue-associated gene sets among differentially expressed genes (DEGs) at each exposure time point (1, 2, 4, 8, 24, and 48 hours; see Table S12). DEGs from each gene set were analyzed using the Preferred tissue (modEncode RNA_seq) gene set. The top three “Preferred tissue” gene sets are displayed according to average *p-*value across the time points. Values in parentheses indicate the NaAsO_2_ dose used in the exposure. The digestive system consistently shows the strongest enrichment across all conditions and time points. (B, C) Heatmaps displaying decreasing log_2_ fold changes in expression for gene families associated with digestion and cholesterol trafficking. Gene expression changes are presented separately for (B) females and (C) males across 6 exposure time points (1, 2, 4, 8, 24, and 48 hours). Gene names in **bold** red exhibit significant down-regulation in both sexes. Gene families include Jonah serine hydrolases (FBgg0001076), lysozymes (FBgg0000606), and Niemann-Pick type C genes (FBhh0000396, FBhh0000397), all of which are significantly downregulated following NaAsO_2_ exposure, indicating disrupted intestinal metabolism.

At the 1.0 mM exposure, many of these DEGs belong to the trypsin- and chymotrypsin-like Jonah family of serine hydrolases, lysozymes, and Niemann-Pick family members involved in cholesterol trafficking—all of which were downregulated following NaAsO_2_ exposure (Figure 6B-C, Table S2-S9). These findings highlight the particular impact of arsenite on digestive and absorption functions in the gut and reveal how tissue transcriptional responses evolve over time: sensing, followed by stress and detoxification responses, and finally adaptive or disease-related changes to exposure.

### Dose-independent responses to NaAsO_2_ encompass genes involved in lipid metabolism, chromatin architecture, and endocrine signaling

While the time-resolved analyses highlighted how transcriptional responses evolve with dose and exposure duration, we next sought to define a set of arsenite-responsive genes that are robustly induced during acute exposure and shared across experimental conditions. To do so, we identified genes that were differentially expressed in arsenite-exposed flies relative to time-matched controls in all four exposure conditions (male and female flies exposed to 0.25 mM or 1.0 mM NaAsO_2_), regardless of the specific time point at which the response occurred. This analysis yielded 100 genes that define a core arsenite-responsive transcriptional program (Table S16). Importantly, this approach does not reflect an average across time points but instead captures genes that respond reproducibly at one or more stages of the acute exposure time course, consistent with their role in initiating or sustaining the arsenite response. The majority of these genes encode proteins with predicted human orthologs (72/100; Table S17), indicating that this core response reflects conserved biological processes rather than condition- or time-specific effects.

Gene set enrichment analysis of these 100 core genes using PANGEA revealed two interconnected networks that together define a canonical arsenite stress response (Figure 7 and Table S18). One network is dominated by heat shock proteins and closely related stress-response genes (Figure 7A), consistent with early activation of the Hsf pathway observed in the time-resolved analysis. The second network is enriched for glutathione S-transferases, cytochrome P450s, and metallothioneins (Figure 7B), reflecting a coordinated detoxification and metal-response program that persists across doses. In addition to these canonical detoxification genes, this network includes genes involved in one-carbon and methionine metabolism (*Gnmt* and *MsrA*) as well as multiple ecdysteroid 22-kinases, an insect-specific enzyme family linked to steroid hormone signaling and xenobiotic responses (Scanlan et al., 2020). Together, these networks define a conserved transcriptional backbone that integrates proteostasis, redox control, and metabolic adaptation in response to arsenite exposure.

**Figure 7.**
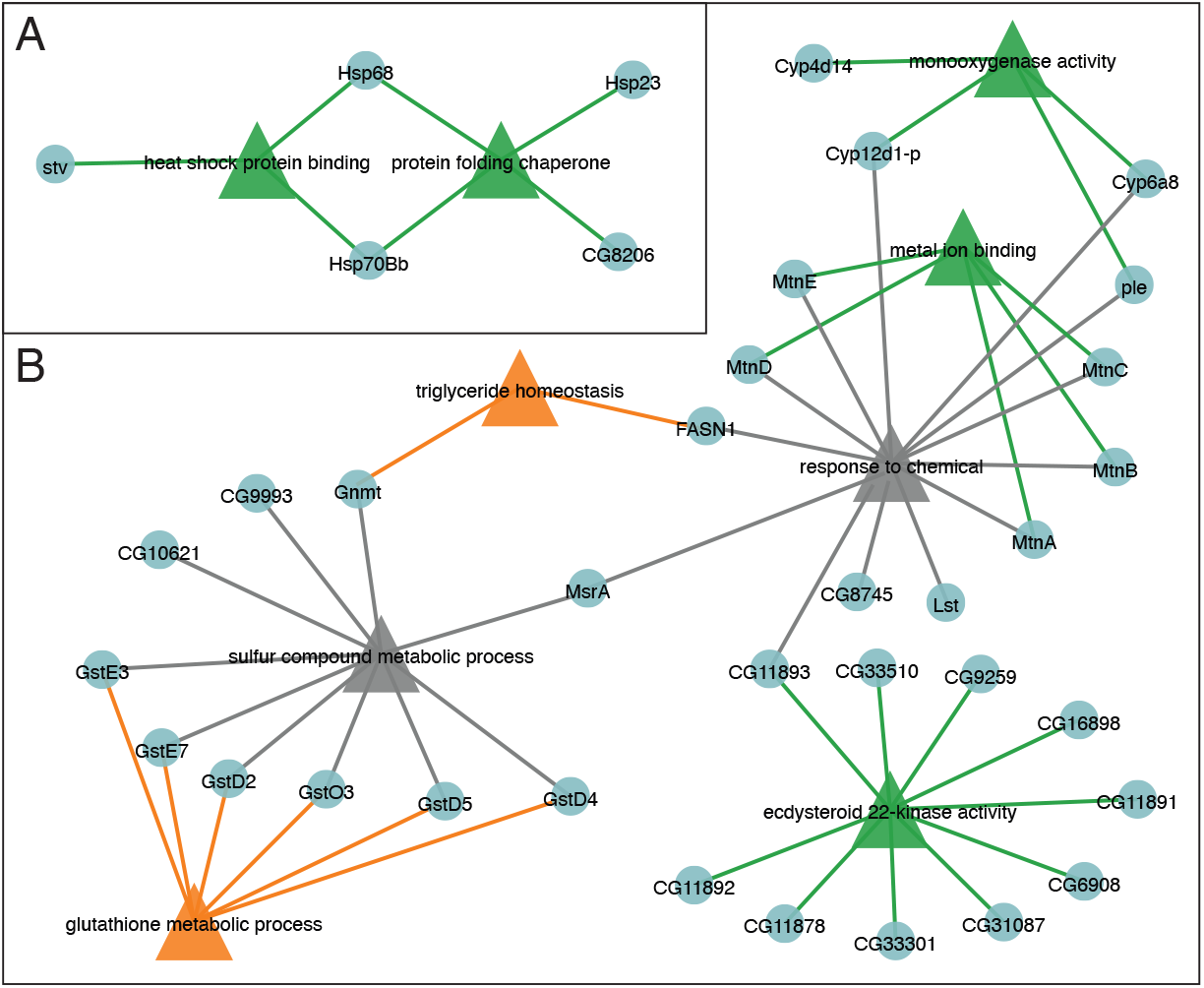
NaAsO_2_ exposure activates a conserved detoxification and stress-response network. The 100 genes consistently differentially expressed across all NaAsO_2_ exposure conditions (see Table S16) were analyzed using PANGEA (Table S18). From these, depicted gene sets were selected based on lowest adjusted *p-*value and minimal overlap between genes and used to generate a gene set node graph. (A,B) The resulting graphs consists of a gene set networks focused on (A) the heat shock response and (B) a series of metabolic pathways that define an integrated defense response to arsenite. Triangle nodes indicate gene sets, and circle nodes denote individual genes.

Beyond the detoxification response, five additional metabolic genes emerged from this analysis (Table S16 and S17). These include *CG33502*, a homolog of Nfu1 involved in Fe-S cluster assembly, and *CG32669*, a member of the SLC5 transporter family – implicating mitochondrial function and Na+ electrochemical gradient in the NaAsO_2_ response. We also observe consistent changes in two lipid metabolism genes: (i) *FASN1* was generally downregulated, (ii) *Dgat2* was upregulated. These findings suggests that arsenite reduces de novo fatty acid synthesis and storage triglycerides, consistent with observations in mice and *C. elegans* (Zdraljevic et al., 2019, Adebayo et al., 2015). Finally, *CG8745*, which encodes an ethanolamine-phosphate phospho-lyase, was also induced in common–a change consistent with elevated phosphoethanolamine levels, a directly testable hypothesis (see metabolism section below).

Two genes with annotated roles in chromatin organization and DNA replication also emerged from this analysis (Tables S16 and S17). We observed consistent induction of *His2B:CG33872*, which encodes a histone H2B variant, and *timeless*, a gene with established roles in circadian regulation and DNA replication fork stability (Vosshall et al., 1994, Sehgal et al., 1994, Vipat and Moiseeva, 2024). Although arsenite exposure has previously been linked to circadian disruption in *Drosophila* (Adebambo et al., 2024), we did not observe coordinated changes in other core clock genes. This suggests that altered *timeless* expression may reflect replication-associated or chromatin-linked stress responses, rather than direct perturbation of circadian timing. However, given that these inferences are based on transcriptional changes alone, additional functional studies will be required to define the precise role of these factors in the arsenite response.

We also identified two genes encoding secreted proteins, *Arc1* and *Limostatin*, that were consistently upregulated across all exposure conditions (Figure 7B and Table S16). *Arc1* has been implicated in lipid metabolism and intercellular communication (Mattaliano et al., 2007, Mosher et al., 2015), while *Limostatin* suppresses insulin secretion in response to nutrient availability (Alfa et al., 2015). The induction of these genes suggests that arsenite exposure engages endocrine and inter-organ signaling pathways, a conclusion that is further supported by the accompanying metabolomic evidence of disrupted carbohydrate metabolism (see below). Together, these findings raise the possibility that arsenite alters hormonal and metabolic signaling early during exposure, although direct effects on endocrine function will require targeted physiological validation.

### NaAsO_2_ exposed flies exhibit transcriptomic profiles indicative of arsenic-associated human disease states

Given that most arsenite-induced DEGs encode proteins conserved in humans, we reanalyzed the RNAseq data using the Phenotype gene set options in PANGEA, which draws on disease annotations from the Alliance of Genome Resources and phenotypes from FlyBase, to determine whether transcriptional changes in flies correspond to gene expression patterns associated with human arsenic-related diseases. This analysis was used to assess whether early transcriptional responses to acute arsenite exposure in flies overlap with gene sets that have been associated with arsenic-related diseases in humans, rather than to infer disease phenotypes directly. Focusing on the 8-hour exposure datasets, we observed significant enrichment of fly DEGs in multiple disease-associated gene sets, including cataracts, renal failure, peripheral vascular disease, hyperlipidemia, pancreatitis, and cardiovascular disorders (Figures 8 and Table S19). Importantly, these disease associations are known to arise from chronic arsenic exposure in humans (Jomova et al., 2011), and their appearance at early time points in flies suggests that arsenite rapidly engages molecular pathways that are implicated in disease risk.

**Figure 8.**
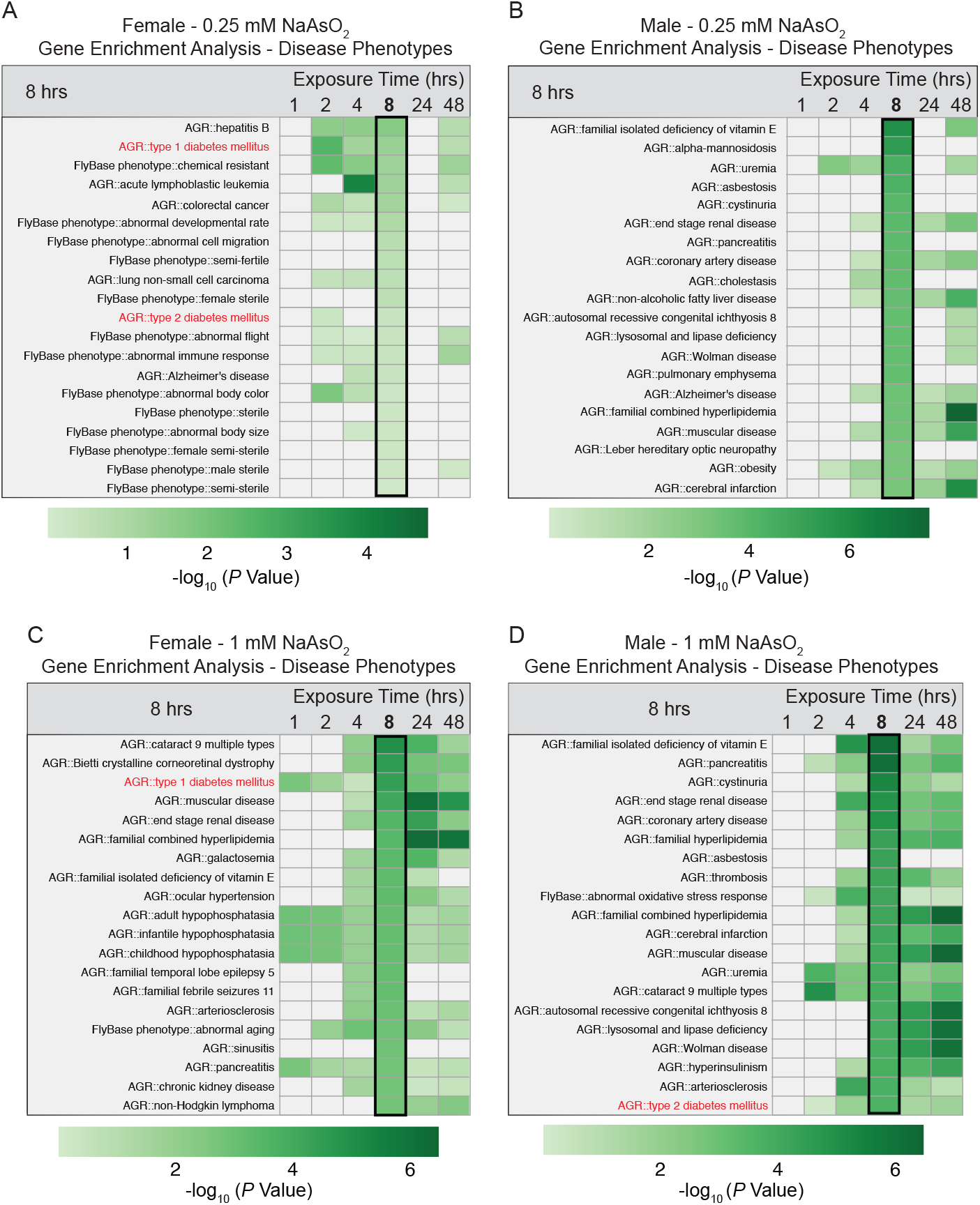
Disease phenotype enrichment among DEGs in female and male *Drosophila melanogaster* exposed to NaAsO_2_. Gene enrichment analysis was performed using PANGEA to identify disease-associated gene sets among differentially expressed genes (DEGs) in female and male flies exposed to either (A, B) 0.25 mM or (C, D) 1.0 mM NaAsO_2_ across a 48-hour time course (see Table S19). Disease annotations were derived from the Alliance of Genome Resources (AGR) and FlyBase phenotype databases. Heatmaps display the top 20 enriched disease phenotypes at the 8 hr time point, rank ordered by significance (indicated by –log_10_ *p*-value), as well as the significance value for those gene sets at all other timepoints. Gene sets associated with Type 1 and 2 diabetes are highlighted in red. The entire analysis for all timepoints can be found in Table S19. Note that high-dose exposure (1mM) resulted in broader and more sustained enrichment of disease phenotypes compared to the 0.25 mM condition.

Although disease associations varied by exposure level and sex, one pattern was consistent: genes linked to diabetes mellitus—both Type 1 and Type 2—were enriched across all datasets (Figure 8 and Table S19]). Notably, Type 1 diabetes–associated genes were significantly enriched in females after just 1 hour of exposure to 1.0 mM NaAsO_2_ (Figure 8 and Table S19), one of the few disease-related sets reaching significance at this early time point (Table S19). This early enrichment coincided with induction of *Limostatin*, a secreted peptide known to suppress insulin secretion in flies (Alfa et al., 2015), which was upregulated across exposure conditions (Table S16).

*Limostatin* is functionally analogous to somatostatin in mammals, a peptide hormone that negatively regulates insulin release during nutrient deprivation (Rorsman and Huising, 2018). Additionally, the insulin receptor (*InR*) was upregulated in males at multiple timepoints (Tables S3, S7), consistent with previous observations that increased *InR* expression correlates with reduced insulin signaling in flies (Puig and Tjian, 2005). Together, these transcriptional changes are consistent with perturbations to endocrine and glucose-regulatory pathways previously implicated in arsenic-associated metabolic dysfunction in human populations (Navas-Acien et al., 2008, Rahimi Kakavandi et al., 2023, Grau-Pérez et al., 2017).

### NaAsO_2_ exposure induces metabolic defects indicative of a diabetic state

Having established that NaAsO_2_ exposure induces rapid transcriptional changes in conserved gene networks, including pathways linked to endocrine signaling and glucose homeostasis, we next turned to the metabolomics data to assess the functional metabolic consequences of these responses. Although earlier analyses indicated that metabolic remodeling follows initial gene expression changes, examining the metabolite profiles in greater detail allowed us to identify which metabolic pathways are most sensitive to arsenite exposure. This analysis revealed an early and pronounced disruption of central carbon metabolism, with pathways governing carbohydrate utilization emerging as a prominent feature of arsenite-induced metabolic stress. Among the top 25 significantly altered metabolites, nearly half were associated with carbohydrate metabolism (Figure 9A–B; Tables S20 and S21). Glucose and lactate levels increased significantly between 4 and 8 hours of exposure (Figure 9A–B; Figure S5A–D; Tables S20 and S21), consistent with early disruption of glycolytic metabolism. Pentose sugars (ribose/ribulose) also accumulated markedly (Figure 9A–B; Tables S20 and S21). These changes were accompanied by elevated levels of methylglyoxal (Figure 9A–B; Figure S5E–F), a reactive byproduct of glycolysis whose accumulation is consistent with increased glycolytic flux (de Bari, 2020). Because lactate and methylglyoxal are established biomarkers of dysregulated glucose metabolism in humans (Moraru et al., 2018, Schalkwijk and Stehouwer, 2020, Lu et al., 2024), these findings support the conclusion that glycolytic dysfunction is a central component of arsenite-induced metabolic stress and align with the transcriptional signatures observed above.

**Figure 9.**
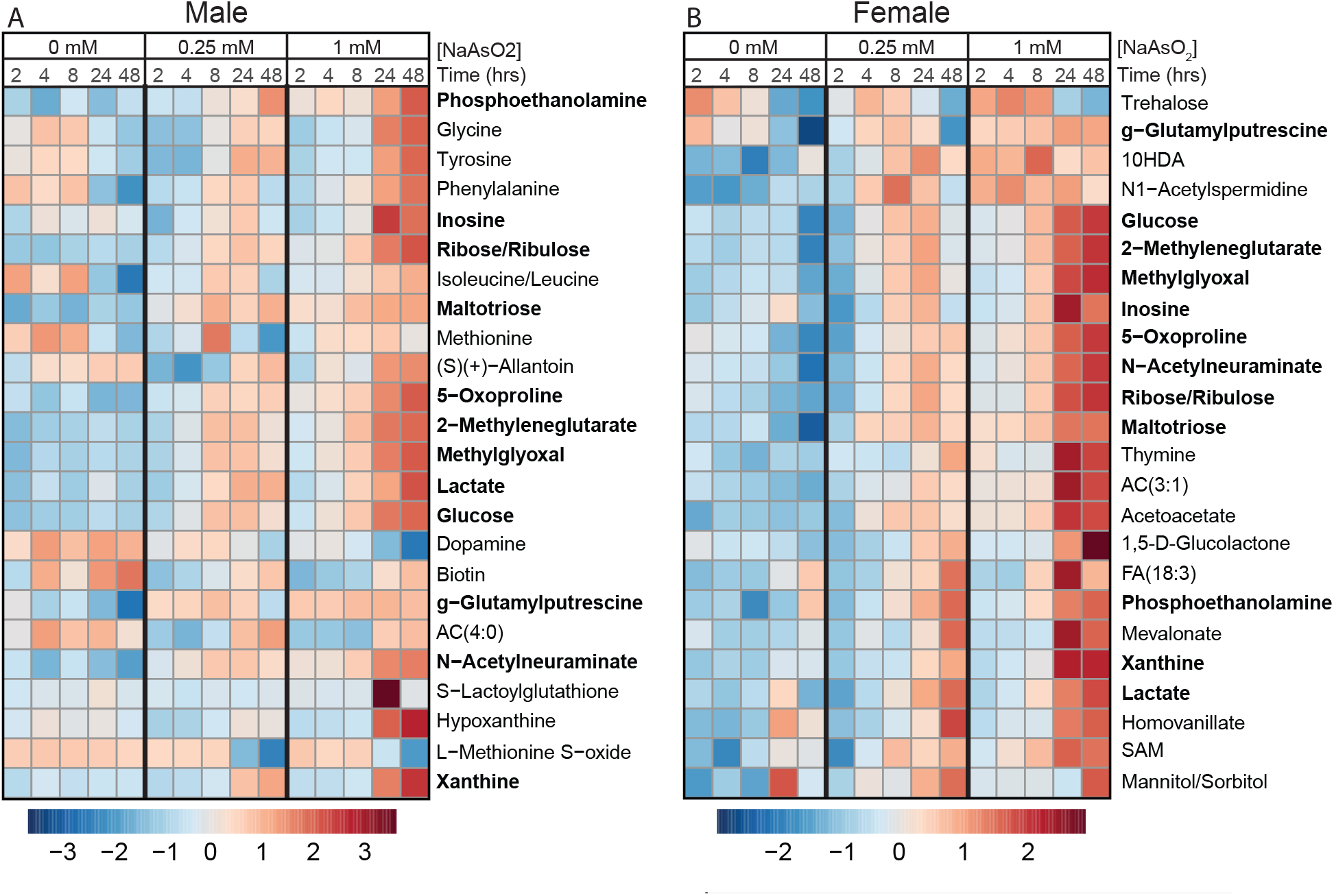
The metabolome of male and female *Drosophila* changes in response to dose and exposure duration. Heatmaps showing top 25 significantly changed metabolites for (A) male and (B) female adult *Drosophila* exposed to both 0.25 mM and 1.0 mM NaAsO_2_ across time (see Tables S10 and S11). The 12 metabolites that are in common between the sexes are highlighted in bold. Each column represents the scaled average metabolite abundance for the six biological replicates analyzed at each time point. Heatmaps were generated using Metaboanalyst 6.0. Statistical significance determined using an ANOVA test. Relative abundance is visualized using Pareto scaling, with elevated levels in red and decreased levels in dark blue.

Beyond carbohydrate metabolism, NaAsO_2_ exposure disrupted several interconnected metabolic pathways associated with oxidative and xenobiotic stress responses. These changes converge on processes that support cellular redox balance and antioxidant capacity, consistent with arsenite-induced oxidative burden.

#### (i) Methionine and folate/one carbon metabolism

Multiple intermediates linked to one-carbon metabolism—including methionine, L-methionine-S-oxide, and S-adenosylmethionine (SAM)—were among the most significantly altered metabolites following NaAsO_2_ exposure (Figure 9A–B; Tables S20 and S21). These metabolites participate in the SAM cycle, which serves as the primary cellular source of methyl groups and contributes to the generation of NADP(H) and glutathione through downstream one-carbon flux (Ducker and Rabinowitz, 2017). Perturbation of this pathway is therefore likely to have broad consequences for redox homeostasis and detoxification capacity. Although the mechanistic basis for these changes remains to be resolved, altered one-carbon metabolism provides a plausible metabolic link between arsenite exposure and increased demand for antioxidant and xenobiotic defense systems.

#### (ii) Oxidative Stress Markers

Consistent with elevated oxidative stress, levels of 5-oxoproline—an established marker of oxidative burden—were increased following arsenite exposure (Figure 9A–B; Tables S20 and S21) (Van Der Pol et al., 2017, Pederzolli et al., 2007, Pederzolli et al., 2010). In addition, cystine levels increased whereas ascorbate levels decreased in flies exposed to 1.0 mM NaAsO_2_, particularly at later time points (8, 24, and 48 hours; Figure 10A–F; Tables S20 and S21). This reciprocal pattern is consistent with disruption of the cystine/cysteine redox cycle and increased utilization of ascorbate as an antioxidant buffer. Together, these changes indicate sustained oxidative stress and suggest a shift toward reliance on non-enzymatic antioxidant systems during prolonged arsenite exposure.

**Figure 10.**
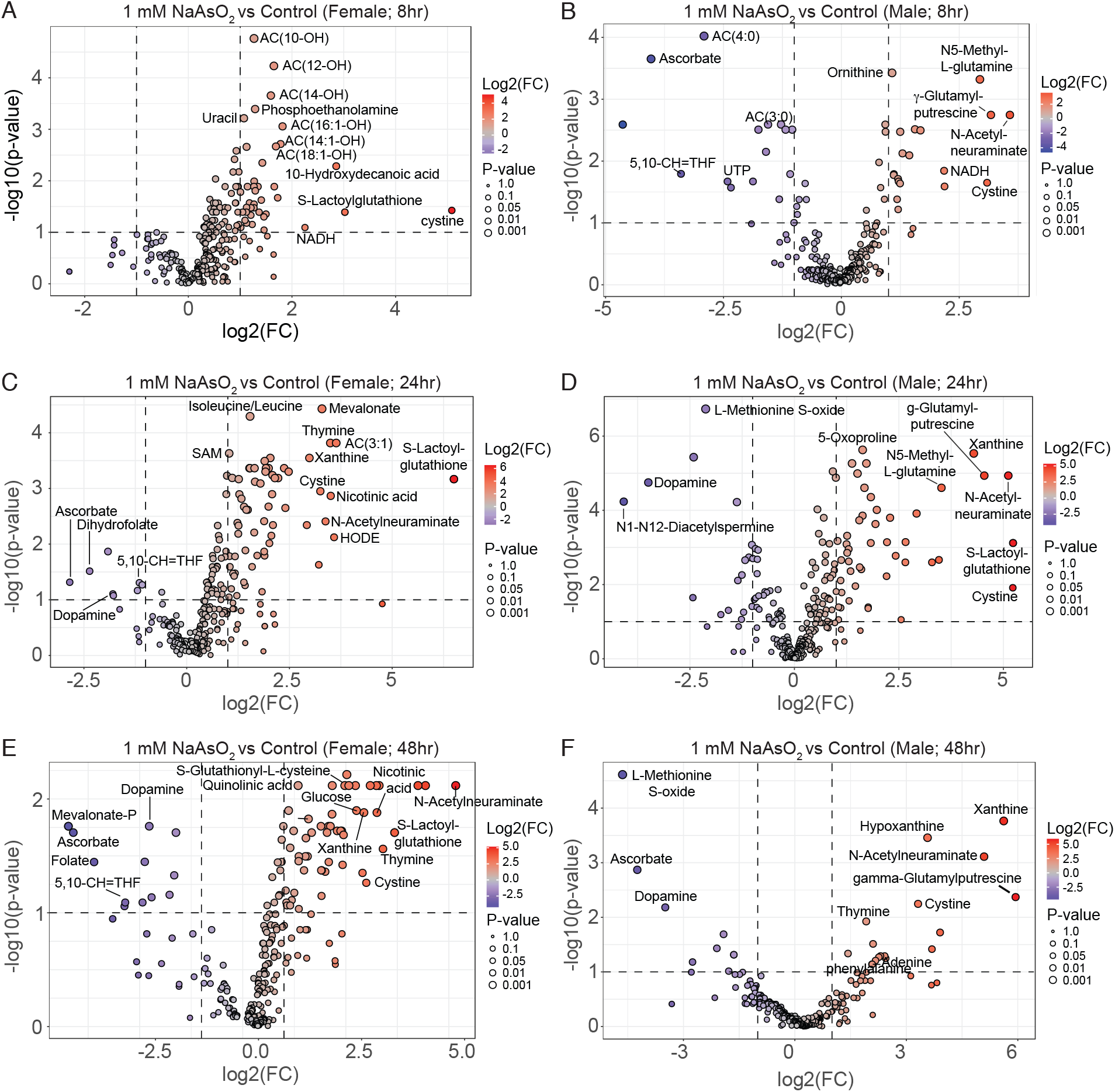
Comparison of metabolite levels between *Drosophila* exposed to 1.0 mM NaAsO_2_ and unexposed controls. Volcano plots comparing relative metabolite levels between NaAsO_2_-treated and untreated flies for the following conditions: (A) Female flies exposed for 8 hours. (B) Male flies exposed for 8 hours. (C) Female flies exposed for 24 hours. (D) Male flies exposed for 24 hours. (E) Female flies exposed for 48 hours. (F) Male flies exposed for 48 hours. For all exposures, relative levels are determined by the average value of n=6 samples each containing 20 flies. Vertical dashed line represents a log_2_ fold change (FC) of 2 and horizontal dashed line represents -log_10_ p-value of 1.

#### (iii) Purine metabolism

Inosine and xanthine levels increased (Figure 9A-B). Xanthine showed particularly large increases at 24 and 48 hours, ranking among the most significantly altered metabolites (Figure 9 and 10C-F and Tables S20 and S21). Importantly, *Drosophila* purine metabolism was previously linked to antioxidant defense, as mutations in the *rosy* gene, which encodes the *Drosophila* homolog of xanthine dehydrogenase, render flies sensitive to oxidative stress (Hilliker et al., 1992). Thus, elevated xanthine levels likely reflect a metabolic response to increased oxidative burden. We also note these changes potentially intersect with folate/one-carbon metabolism, as purine biosynthesis requires two one-carbon units from 10-formyl-tetrahydrofolate (Ducker and Rabinowitz, 2017).

In addition to the metabolic changes associated with oxidative stress, phosphoethanolamine was consistently elevated in arsenite-exposed flies across doses and time points (Figure 9A–B; Tables S20 and S21). This observation is notable because ***CG8745***, which is predicted to encode an ethanolamine-phosphate phospho-lyase, is among the 100 genes induced under all exposure conditions (Tables S16 and S17). Although the functional consequences of these changes remain to be determined, the coordinated elevation of phosphoethanolamine and induction of a gene linked to its catabolism suggests that phosphoethanolamine metabolism responds robustly to arsenite exposure and may represent a sensitive indicator of arsenite-induced metabolic stress.

Taken together, the metabolomic data indicate that NaAsO_2_ exposure perturbs central carbon metabolism, elevates reactive metabolic byproducts, and engages oxidative stress–associated pathways. When considered alongside the transcriptional changes in genes involved in endocrine and metabolic regulation, these findings support the use of *Drosophila* as a tractable whole-organism system for dissecting conserved metabolic responses to arsenic exposure and for identifying early molecular features associated with arsenite-induced metabolic dysfunction.

## DISCUSSION

This study provides a comprehensive, time-resolved multi-omic analysis of arsenite exposure in *Drosophila melanogaster*, revealing how molecular responses evolve from the earliest stages of exposure through the onset of lethality. Unlike previous studies that focused on individual genes, isolated pathways, or single exposure time points (Muñiz Ortiz et al., 2009, Chaturvedi et al., 2025), our integrated transcriptomic and metabolomic approach resolves the temporal ordering of molecular responses across dose and time. This resolution enables identification of early, coordinated activation of conserved stress, detoxification, endocrine, and metabolic programs that precede large-scale metabolic remodeling. Consistent with this early pathway engagement, disease-associated gene set enrichment analyses indicate that acute arsenite exposure in flies engages molecular pathways implicated in arsenic-associated chronic diseases in humans. Together, these findings highlight the translational utility of the fly model for identifying early molecular signatures and initiating events relevant to arsenic-associated disease risk.

Importantly, among the 100 genes consistently induced across all exposure conditions, 72 encode proteins with predicted human orthologs (Table S17), indicating that the core arsenite-responsive transcriptional program in *Drosophila* is largely evolutionarily conserved. Notably, this level of conservation is consistent with previous observations that ∼70% of adversity- and disease-linked genes are shared among distantly related species (Colbourne et al., 2022, Wangler et al., 2017). Rather than defining discrete regulatory networks, these conserved genes converge on a small number of recurring functional themes, including oxidative stress defense, xenobiotic metabolism, and endocrine-linked metabolic regulation—processes that are broadly implicated in arsenic toxicity in humans. For example, multiple glutathione S-transferases, cytochrome P450 enzymes, and metallothioneins were consistently induced, reflecting a conserved detoxification and redox-buffering response to arsenite across species. In addition, conserved regulators of metabolic signaling, including *Limostatin* and the insulin receptor (*InR*), were consistently altered, suggesting that arsenite exposure engages endocrine pathways linked to glucose regulation early in the response. Together, these findings support the use of *Drosophila* as a whole-organism model for identifying conserved early molecular responses and biomarkers associated with arsenic exposure. In this context, arsenic serves as a well-characterized exemplar, illustrating how time-resolved multi-omic analysis in *Drosophila* can reveal conserved early molecular responses relevant to environmental health.

Both RNA-seq and semi-targeted metabolomic analyses revealed coordinated molecular responses to NaAsO_2_ exposure, with particularly strong disruption of carbohydrate metabolism. Within hours of exposure, levels of glucose, lactate, and methylglyoxal increased markedly, coincident with transcriptional changes in endocrine regulators such as *Limostatin*, a peptide that suppresses insulin secretion in flies (Alfa et al., 2015). Similar alterations in insulin signaling and glycolytic control have been associated with chronic arsenic exposure in human populations (Navas-Acien et al., 2008, Lai et al., 1994, Grau-Pérez et al., 2017). The rapid onset of these metabolic changes highlights the sensitivity of carbohydrate metabolism to arsenite exposure and supports the existence of a conserved axis linking arsenic toxicity to metabolic dysregulation.

Another key advance from this work is the temporal ordering of arsenite-responsive transcriptional programs. Our data suggest that the coordinate activation of three transcription factors drives many of the initial responses. First, we observed rapid induction of heat-shock protein genes, which are canonical targets of the Heat Shock Factor (Hsf) family of transcription factors (Wu et al., 1987, Clos et al., 1990). This result is consistent with the known role of Hsf in protecting cells against proteotoxic insults from a variety of environmental stresses, including heavy metals and toxins (Akerfelt et al., 2010). This initial response associated with Hsf is followed by upregulation of Phase I xenobiotic and oxidative stress response genes, including glutathione S-transferases and cytochrome P450s—targets of Cap ‘n’ collar (Cnc), the *Drosophila* homolog of Nrf2 (Sykiotis and Bohmann, 2008, Misra et al., 2013). Similar to mammalian Nrf2, Cnc is controlled by its cytoplasmic inhibitor Keap1, which suppresses Nrf2 transcriptional activity and targets this transcription factor for degradation by the proteasome (Baird and Yamamoto, 2020, Sykiotis and Bohmann, 2008). Oxidative stress and exposure to electrophiles disrupts sulfhydryl bonds within Keap1, releasing Cnc/Nrf2 and activating the downstream transcriptional program (Baird and Yamamoto, 2020). In this regard, studies from human cells demonstrated that inorganic arsenic indeed disrupts critical sulfhydryl bonds within Keap1, resulting in Nrf2 activation (Lau et al., 2013). Thus, arsenite potentially activates this coordinated transcriptional network directly in the fly by changing the structure of Keap1, which requires no transcriptional or translational control. Finally, the early transcriptional responses driven by Hsf and Cnc/Nrf2 occur in parallel with induction of five *Drosophila* metallothionein genes. Increased expression of these genes likely reflects activation of the MTF-1 transcription factor, which is known to drive *Mtn* gene expression in response to heavy metals (Zhang et al., 2001).

The early activation of Hsf target genes relative to Cnc/Nrf2 transcriptional networks indicates that the heat shock response represents one of the earliest cellular defenses engaged following arsenite exposure. This temporal ordering is notable given that arsenic is known to directly disrupt Keap1 function and activate Cnc/Nrf2 signaling, yet our time-resolved data reveal that Hsf-dependent transcriptional programs are mobilized more rapidly. One explanation for this pattern is that arsenite rapidly induces widespread proteotoxic stress (Jacobson et al., 2012, Guerra-Moreno et al., 2015), leading to protein misfolding and increased demand for molecular chaperones. Under such conditions, pre-existing pools of heat shock proteins become engaged in stabilizing damaged or misfolded proteins, thus relieving chaperone-mediated repression of Hsf and permitting its rapid activation (Akerfelt et al., 2010, Morimoto, 2011). This mechanism is consistent with established models in which Hsf functions as a sensor of proteostasis imbalance, responding swiftly to disruptions in protein folding homeostasis. In this framework, Hsf-driven transcription represents an early response to proteostasis disruption, while activation of Cnc/Nrf2-dependent detoxification and redox pathways reflects a coordinated but temporally distinct response to arsenite-induced oxidative and electrophilic stress.

Our time-resolved analysis also suggests temporal separation between transcriptional and metabolic responses to arsenite. Principal component analysis of the RNA-seq data shows that gene expression changes emerge rapidly, with distinct exposure clustering by dose as early as 8 hours. In contrast, clustering analysis of the metabolomic profiles reveals segregation only at later time points (Figures 1–2). This observation, however, does not imply that metabolic activity remains static at the early stages of arsenite exposure. Flux through a pathway can increase substantially without causing large shifts in steady-state metabolite concentrations if production and consumption rates rise in parallel. In the context of arsenite exposure, enhanced detoxification and antioxidant demands likely accelerate flux through folate metabolism, the methionine cycle, and glutathione biosynthesis, while homeostatic mechanisms buffer pool sizes to prevent harmful depletion or imbalance. This dynamic could explain why strong transcriptional enrichment of genes involved in glutathione metabolism coincides with relatively modest changes in relevant metabolite levels, particularly at early time points. In contrast, the accumulation of xanthine and other purine metabolites at later stages potentially reflects significant, destabilizing imbalances in cellular metabolism that occur once homeostatic mechanisms are no longer sufficient to buffer stress. Such unobserved regulation underscores the importance of integrating flux-based approaches with transcriptomic and metabolomic profiling in future studies.

While our multi-omic approach highlights the dynamic nature of the arsenic defense response, it also reveals that molecular pathways linked to chronic arsenic-associated diseases are engaged at the earliest stages of acute exposure. This observation underscores the importance of conducting whole-organism, time-resolved analyses to identify molecular initiating events that precede overt pathology and highlights a future need for tissue-specific studies that refine the mechanistic interpretation of these early molecular signatures. For example, gene set enrichment analysis suggests a key role for intestinal physiology and barrier function in the response to arsenite, pointing to the need for tissue-specific and single-cell analyses to uncover organ-level contributions. In this context, recent single-nucleus RNA-seq profiling of the *Drosophila* brain following NaAsO_2_ exposure complements our analysis and reveals both similarities and differences in cell-specific arsenite responses (Chaturvedi et al., 2025). Notably, seven genes from our core signature (*MtnA, MsrA, CG8745, tim, ple, Arc1*, and *CG12224*) were also significantly altered in the brain dataset (Chaturvedi et al., 2025). Moreover, both datasets converge on the same detoxification gene families, including glutathione S-transferases, cytochrome P450s, UDP-glucuronosyltransferases, metallothioneins, and methionine sulfoxide reductases (Chaturvedi et al., 2025).

Taken together, our findings establish *Drosophila* as a uniquely powerful whole-organism model for investigating the molecular initiating events that precede onset of toxicant-induced diseases. By leveraging its genetic tractability, rapid life cycle, multiple organ systems, and capacity for sex-specific analysis, we demonstrate how the fly can illuminate evolutionarily conserved biological responses and uncover early biomarkers of adverse outcome pathways (Ankley and Edwards, 2018). As toxicology continues to evolve toward mechanistic, predictive, and ethically responsible approaches, *Drosophila* is poised to play a central role in bridging experimental discovery with human health relevance (Rand et al., 2023). Our study not only advances the application of multi-omic strategies in NAMs but also lays the groundwork for future efforts to integrate fly-based insights into regulatory frameworks and public health initiatives aimed at mitigating the global burden of environmental toxicants such as arsenic.

## SUPPLEMENTAL FIGURE LEGENDS

**Figure S1. Dose-response curves of *Drosophila* adult males and females exposed to NaAsO**_**2**_. Dose-response curves illustrate the measured proportions of dead (A) female and (B) male Oregon-R flies at increasing concentrations of NaAsO_2_ following a 48 hr exposure. Both the best fit line and the LD_10_, LD_25_, and LD_50_ were calculated using a Dichotomous Hill Model. n=6 vials at each concentration with 20 flies per vial. (A,B – see text in lower right of panel) Male and female dose response curves were statistically analyzed with a global nonlinear regression model analysis using an extra sum-of-squares F test. This analysis revealed a significant difference in the LD_25_ and LD_50_ values between Oregon-R male and female flies (LD_10_ F-statistic of F(1,112) = 1.581, *p* = 0.2112; LD_25_ F-statistic of F(1,112) = 5.586, *p* = 0.0198; LD_50_ F-statistic of F(1,112) = 16.85, *p* < 0.0001). However, the hillslope of the dose response curve was similar between males and females (F-statistic of F(1,112) = 0.3659, *p* = 0.5465).

**Figure S2. A comparison of DEG in males and females exposed to 1.0 mM NaAsO**_**2**_. Venn diagrams illustrating the overlap between up- and down-regulated DEGs in males and females at 4, 8, 24, and 48 hrs. Note that less than 50% of the DEGs at any given time point are shared between the two sexes. The 1 hr and 2 hr time points were excluded from this analysis due to the low number of DEGs.

**Figure S3. Gene Set Enrichment Analysis of DEGs in female *Drosophila melanogaster* exposed to 0.25 mM NaAsO**_**2**_ **across a 48-hour time course**.

As in Figure 3, PANGEA was used to identify significantly overrepresented gene sets among differentially expressed genes (DEGs) at each exposure time point (see Methods and Table S3). DEGs from each time point were simultaneously analyzed for enrichment using four annotation sources: (i) SLIM2 GO Biological Process, (ii) FlyBase signaling pathways (experimental evidence), (iii) KEGG Pathway (*D. melanogaster*), and (iv) REACTOME Pathway (*D. melanogaster*). The top 10 enriched gene sets for each time point are presented with heatmaps of their rank-ordered adjusted *p*-value at the corresponding time (outlined heat map column), along with their significance (green gradient) or lack thereof (grey) at the other time points.

**Figure S4. Gene Set Enrichment Analysis of DEGs in male *Drosophila melanogaster* exposed to 0.25 mM NaAsO**_**2**_ **across a 48-hour time course**.

As in Figure 3, PANGEA was used to identify significantly overrepresented gene sets among differentially expressed genes (DEGs) at each exposure time point (see Methods and Table S5). The top 10 enriched gene sets for each time point are presented with heatmaps of their rank-ordered adjusted *p*-value at the corresponding time (outlined heat map column), along with their significance (green) or lack thereof (grey) at the other time points.

**Figure S5. NaAsO**_**2**_ **exposure disrupts glycolytic metabolism in a dose- and time-dependent manner**. Box and whiskers plots displaying normalized levels of (A, B) glucose, (C, D) lactate, and (E, F) methylglyoxal in adult flies exposed to 0 mM (control), 0.25 mM, or 1.0 mM NaAsO_2_ over a 48-hour time course. Analysis conducted with Metaboanalyst 6.0. Data were normalized to sample mass and processed with Log Transformation and Pareto scaling. Statistical analysis was conducted using ANOVA with FDR correction (FDR < 0.05), followed by Fisher’s Least Significant Difference post hoc test (see Tables S20 and S21). *\*p* < 0.05 relative to untreated time-matched control. n=6 samples per condition.

## SUPPLEMENTAL TABLES

**Table S1**. Data used to generate dose response curves in Figure s1. n=6 exposure vials per NaAsO_2_ concentration; 20 animals per vial.

**Table S2**. RNA-seq results comparing gene expression between Oregon-R Males exposed to NaAsO_2_. Adult males were exposed to 0 mM and 0.25 mM NaAsO_2_ for 1, 2, 4, 8, 24 or 48 hours prior to collection.

**Table S3**. RNA-seq results comparing gene expression between Oregon-R Males exposed to NaAsO_2_. Adult males were exposed to 0 mM and 0.25 mM NaAsO_2_ for 1, 2, 4, 8, 24 or 48 hours prior to collection. Only genes displaying significant changes in gene expression (log_2_ fold change ≥ |1| and an adjusted *p-*value of ≤0.05) are included in these tables.

**Table S4**. RNA-seq results comparing gene expression between Oregon-R females exposed to NaAsO_2_. Adult females were exposed to 0 mM and 0.25 mM NaAsO_2_ for 1, 2, 4, 8, 24 or 48 hours prior to collection.

**Table S5**. RNA-seq results comparing gene expression between Oregon-R females exposed to NaAsO_2_. Adult females were exposed to 0 mM and 0.25 mM NaAsO_2_ for 1, 2, 4, 8, 24 or 48 hours prior to collection. Only genes displaying significant changes in gene expression (log_2_ fold change ≥ |1| and an adjusted *p-*value of ≤0.05) are included in these tables.

**Table S6**. RNA-seq analysis comparing gene expression between groups of Oregon-R males exposed to NaAsO_2_. Adult males were exposed to 0 mM and 1.0 mM NaAsO_2_ for 1, 2, 4, 8, 24 or 48 hours prior to collection.

**Table S7**. RNA-seq analysis comparing gene expression between groups of Oregon-R males exposed to NaAsO_2_. Adult males were exposed to 0 mM and 1.0 mM NaAsO_2_ for 1, 2, 4, 8, 24 or 48 hours prior to collection. Only genes displaying significant changes in gene expression (log_2_ fold change ≥ |1| and an adjusted *p-*value of ≤0.05) are included in these tables.

**Table S8**. RNA-seq analysis comparing gene expression between groups of Oregon-R females exposed to NaAsO_2_. Adult females were exposed to 0 mM and 1.0 mM NaAsO_2_ for 1, 2, 4, 8, 24 or 48 hours prior to collection.

**Table S9**. RNA-seq analysis comparing gene expression between groups of Oregon-R females exposed to NaAsO_2_. Adult females were exposed to 0 mM and 1.0 mM NaAsO_2_ for 1, 2, 4, 8, 24 or 48 hours prior to collection. Only genes displaying significant changes in gene expression (log_2_ fold change ≥ |1| and an adjusted *p-*value of ≤0.05) are included in these tables.

**Table S10**. Metabolomic analysis of Oregon-R Males exposed to NaAsO_2_. Adult Males were exposed to 0 mM, 0.25 mM, and 1.0 mM NaAsO_2_ for 2, 4, 8, 24, or 48 hours prior to collection. n=6 samples per condition; 20 adult males per sample. Data represented as ion counts normalized to sample mass.

**Table S11**. Metabolomic analysis of Oregon-R females exposed to NaAsO_2_. Adult females were exposed to 0 mM, 0.25 mM, and 1.0 mM NaAsO_2_ for 2, 4, 8, 24, or 48 hours prior to collection. n=6 samples per condition; 20 adult females per sample. Data represented as ion counts normalized to sample mass.

**Table S12**. The number of differentially expressed genes (DEGs) in NaAsO2-exposed adult male and female flies sorted by dose and time point. The tabulated DEGs were identified as having a log2 fold change ≥ |1| and an adjusted p-value of ≤0.05. The full list of DEGs can be found in Table S3, S5, S7, and S9.

**Table S13**. The number of significantly altered metabolites in NaAsO2-exposed adult male and female flies sorted by dose and time point. The tabulated metabolites were identified as having a log2 fold change ≥ |1| and an adjusted p-value of ≤0.01. The full list of metabolites can be found in Table S10 and S11.

**Table S14**. PANGEA analysis of differentially expressed genes in flies exposed to either 0.25 mM or 1.0 mM NaAsO_2_ for 1,2,4,8,24, and 48 hrs. DEGs from each time point were simultaneously analyzed for enrichment using four annotation sources: (i) SLIM2 GO Biological Process, (ii) FlyBase signaling pathways (experimental evidence), (iii) KEGG Pathway (*D. melanogaster*), and (iv) REACTOME Pathway (*D. melanogaster*). **Table S15**. PANGEA analysis of differentially expressed genes in male and female flies exposed to either 0.25 mM or 1.0 mM NaAsO_2_ for 1,2,4,8,24, and 48 hrs. DEGs from each time point were simultaneously analyzed for enrichment using the Preferred tissue (modEncode RNA_seq) gene set.

**Table S16. Core transcriptional response to NaAsO**_**2**_ **exposure shared across sexes and doses**. List of 100 genes consistently differentially expressed in adult *Drosophila melanogaster* under all four exposure conditions (male and female flies at 0.25 mM and 1.0 mM NaAsO_2_). Genes are annotated with FlyBase identifiers.

**Table S17. Human orthologs of conserved arsenite-responsive genes identified by DIOPT analysis**. Predicted human orthologs for the 100 *Drosophila melanogaster* genes consistently differentially expressed across all NaAsO_2_ exposure conditions (see Table S16). Orthology was assessed using the Drosophila RNAi Screening Center Integrative Ortholog Prediction Tool (DIOPT). High-confidence orthologs (DIOPT score ≥10) highlight conserved pathways involved in detoxification, stress response, metabolism, chromatin architecture, and endocrine signaling.

**Table S18**. PANGEA analysis of 100 genes consistently differentially expressed in adult *Drosophila melanogaster* under all four exposure conditions (male and female flies at 0.25 mM and 1.0 mM NaAsO_2_).

**Table S19. Disease phenotype enrichment among DEGs in female and male *Drosophila melanogaster* exposed to NaAsO**_**2**_. Gene enrichment analysis was performed using PANGEA to identify disease-associated gene sets among differentially expressed genes (DEGs) in female and male flies exposed to either 0.25 mM or 1.0 mM NaAsO_2_ across a 48-hour time course.

**Table S20. Statistical analysis of male metabolomics data**. Metaboanalyst 6.0 was used to analyze data in Table S12. Statistical analysis conducted using ANOVA followed by a post hoc Fisher’s Least Significant Difference test.

**Table S21. Statistical analysis of female metabolomics data**. Metaboanalyst 6.0 was used to analyze data in Table S11. Statistical analysis conducted using ANOVA followed by a post hoc Fisher’s Least Significant Difference test.

## ACKNOWLEDGMENTS

We thank the following individuals for helping with the testing and optimization of the exposure protocol: Marilyn Clark, Emma Rose Gallant, Morgan Marsh, Kyle McClung, Andy Puga, and Cameron Stockbridge. The authors would like to thank the Indiana University Bloomington Drosophila Stock Center (BDSC) for fly stocks. RNA sequencing was conducted at the Indiana University Center for Genomics and Bioinformatics. We also are indebted to Claire Hu and the Perrimon lab who in collaboration with GO and FlyBase curators and developers, created the PANGEA enrichment tool. Much of our analysis would have been more difficult without it. This project received funding from the European Union’s Horizon 2020 Research and Innovation program under Grant Agreement No. 965406. The work presented in this publication was performed as part of the ASPIS Cluster. This output reflects only the authors’ views, and the European Union cannot be held responsible for any use that may be made of the information contained herein. The research conducted by Brian Oliver was supported by the Intramural Research Program of the National Institute of Diabetes and Digestive and Kidney Diseases (NIDDK) within the National Institutes of Health (NIH). The contributions of the NIH author(s) are considered Works of the United States Government. The findings and conclusions presented in this paper are those of the author(s) and do not necessarily reflect the views of the NIH or the U.S. Department of Health and Human Services. This publication was also made possible with support from the Indiana Clinical and Translational Sciences Institute, which is funded in part by Award Number UL1TR002529 from the National Institutes of Health, National Center for Advancing Translational Sciences, Clinical and Translational Sciences Award. The content is solely the responsibility of the authors and does not necessarily represent the official views of the National Institutes of Health. Portions of this project were supported by funds from Indiana University awarded to JRS and the PhyloTox consortium.

## Notes

https://www.ncbi.nlm.nih.gov/geo/query/acc.cgi?acc=GSE241663

https://10.5281/zenodo.18381265

### Competing Interest Statement

The authors have declared no competing interest.

## REFERENCES

Adebambo, T. H., Flores, M. F. M., Zhang, S. L. & Lerit, D. 2024. Arsenic impairs Drosophila neural stem cell mitotic progression and sleep behavior in a tauopathy model. bioRxiv, 2024.08.05.606375.

Adebayo, A. O., Zandbergen, F., Kozul-Horvath, C. D., Gruppuso, P. A. & Hamilton, J. W. 2015. Chronic exposure to low-dose arsenic modulates lipogenic gene expression in mice. J Biochem Mol Toxicol, 29, 1–9.

Agency, U. S. E. P. 2023. Benchmark Dose Software (BMDS). 3.3.2 ed.

Akerfelt, M., Morimoto, R. I. & Sistonen, L. 2010. Heat shock factors: integrators of cell stress, development and lifespan. Nat Rev Mol Cell Biol, 11, 545–55.

Alfa, R. W., Park, S., Skelly, K. R., Poffenberger, G., Jain, N., Gu, X., Kockel, L., Wang, J., Liu, Y., Powers, A. C. & Kim, S. K. 2015. Suppression of insulin production and secretion by a decretin hormone. Cell Metab, 21, 323–334.

Andrews, S. 2010. FastQC: a quality control tool for high throughput sequence data. Cambridge, United Kingdom.

Ankley, G. T. & Edwards, S. W. 2018. The Adverse Outcome Pathway: A Multifaceted Framework Supporting 21(st) Century Toxicology. Curr Opin Toxicol, 9, 1–7.

Anushree Ali, M. Z., Bilgrami, A. L. & Ahsan, J. 2023. Acute exposure to arsenic affects pupal development and neurological functions in drosophila melanogaster. Toxics, 11, 327.

Anushree, A., Ali, Z. & Ahsan, J. 2022. Acute exposure to arsenic affects cognition in drosophila melanogaster larvae. Entomology and Applied Science Letters, 9, 70–78.

Ayotte, J. D., Medalie, L., Qi, S. L., Backer, L. C. & Nolan, B. T. 2017. Estimating the High-Arsenic Domestic-Well Population in the Conterminous United States. Environmental Science & Technology, 51, 12443–12454.

Bahadorani, S. & Hilliker, A. J. 2009. Biological and behavioral effects of heavy metals in Drosophila melanogaster adults and larvae. Journal of insect behavior, 22, 399–411.

Baird, L. & Yamamoto, M. 2020. The Molecular Mechanisms Regulating the KEAP1-NRF2 Pathway. Mol Cell Biol, 40.

Chaturvedi, A., Shankar, V., Simkhada, B., Lyman, R. A., Freymuth, P., Howansky, E., Collins, K. M., Mackay, T. F. C. & Anholt, R. R. H. 2025. Arsenic toxicity in the Drosophila brain at single cell resolution. Front Toxicol, 7, 1636431.

Chowdhury, U. K., Biswas, B. K., Chowdhury, T. R., Samanta, G., Mandal, B. K., Basu, G. C., Chanda, C. R., Lodh, D., Saha, K. C., Mukherjee, S. K., Roy, S., Kabir, S., Quamruzzaman, Q. & Chakraborti, D. 2000. Groundwater arsenic contamination in Bangladesh and West Bengal, India. Environ Health Perspect, 108, 393–7.

Clos, J., Westwood, J. T., Becker, P. B., Wilson, S., Lambert, K. & Wu, C. 1990. Molecular cloning and expression of a hexameric Drosophila heat shock factor subject to negative regulation. Cell, 63, 1085–97.

Colbourne, J. K., Shaw, J. R., Sostare, E., Rivetti, C., Derelle, R., Barnett, R., Campos, B., Lalone, C., Viant, M. R. & Hodges, G. 2022. Toxicity by descent: A comparative approach for chemical hazard assessment. Environmental Advances, 9, 100287.

Ducker, G. S. & Rabinowitz, J. D. 2017. One-Carbon Metabolism in Health and Disease. Cell Metab, 25, 27–42.

Ewels, P., Magnusson, M., Lundin, S. & Käller, M. 2016. MultiQC: summarize analysis results for multiple tools and samples in a single report. Bioinformatics, 32, 3047–3048.

Goldstein, S. & Babich, H. 1989. Differential effects of arsenite and arsenate to Drosophila melanogaster in a combined adult/developmental toxicity assay. Bull. Environ. Contam. Toxicol.;(United States), 42.

Grau-Pérez, M., Kuo, C. C., Spratlen, M., Thayer, K. A., Mendez, M. A., Hamman, R. F., Dabelea, D., Adgate, J. L., Knowler, W. C., Bell, R. A., Miller, F. W., Liese, A. D., Zhang, C., Douillet, C., Drobná, Z., Mayer-Davis, E. J., Styblo, M. & Navas-Acien, A. 2017. The Association of Arsenic Exposure and Metabolism With Type 1 and Type 2 Diabetes in Youth: The SEARCH Case-Control Study. Diabetes Care, 40, 46–53.

Guerra-Moreno, A., Isasa, M., Bhanu, M. K., Waterman, D. P., Eapen, V. V., Gygi, S. P. & Hanna, J. 2015. Proteomic Analysis Identifies Ribosome Reduction as an Effective Proteotoxic Stress Response. J Biol Chem, 290, 29695–706.

Hayot, G., Lloyd, G. R., Diwan, G. D., Keith, N., Smoot, S. R., CRAMER Von Clausbruch, C. A., Kaufman, T. C., Barnard, M., Tindall, A. J., Glaholt, S. P., Massei, R., Martínez, R., Strähle, U., Orsini, L., Russell, R. B., Tennessen, J. M., Scholz, S., Shaw, J. R., Freedman, J. H., Colbourne, J. K., Weiss, C. & Dickmeis, T. 2025. Alternative Vertebrate and Invertebrate Model Organisms Show Similar Sensitivity as Rodents to a Diverse Set of Chemicals. Environ Sci Technol, 59, 25634–25648.

Hilliker, A. J., Duyf, B., Evans, D. & Phillips, J. P. 1992. Urate-null rosy mutants of Drosophila melanogaster are hypersensitive to oxygen stress. Proceedings of the National Academy of Sciences, 89, 4343–4347.

Holsopple, J. M., Smoot, S. R., Popodi, E. M., Colbourne, J. K., Shaw, J. R., Oliver, B., Kaufman, T. C. & Tennessen, J. M. 2023. Assessment of chemical toxicity in adult Drosophila melanogaster. JoVE (Journal of Visualized Experiments), e65029.

Hu, Y., Comjean, A., Attrill, H., Antonazzo, G., Thurmond, J., Chen, W., Li, F., Chao, T., Mohr, S. E. & Brown, N. H. 2023. PANGEA: a new gene set enrichment tool for Drosophila and common research organisms. Nucleic Acids Research, 51, W419–W426.

Hughes, M. F. 2002. Arsenic toxicity and potential mechanisms of action. Toxicology letters, 133, 1–16.

Ivy, N., Mukherjee, T., Bhattacharya, S., Ghosh, A. & Sharma, P. 2023. Arsenic contamination in groundwater and food chain with mitigation options in Bengal delta with special reference to Bangladesh. Environmental Geochemistry and Health, 45, 1261–1287.

Jacobson, T., Navarrete, C., Sharma, S. K., Sideri, T. C., Ibstedt, S., Priya, S., Grant, C. M., Christen, P., Goloubinoff, P. & Tamás, M. J. 2012. Arsenite interferes with protein folding and triggers formation of protein aggregates in yeast. Journal of Cell Science, 125, 5073–5083.

Jomova, K., Jenisova, Z., Feszterova, M., Baros, S., Liska, J., Hudecova, D., Rhodes, C. J. & Valko, M. 2011. Arsenic: toxicity, oxidative stress and human disease. Journal of applied toxicology, 31, 95–107.

Lai, M.-S., Hsueh, Y.-M., Chen, C.-J., Shyu, M.-P., Chen, S.-Y., Kuo, T.-L., Wu, M.-M. & Tai, T.-Y. 1994. Ingested inorganic arsenic and prevalence of diabetes mellitus. American Journal of Epidemiology, 139, 484–492.

Lau, A., Whitman, S. A., Jaramillo, M. C. & Zhang, D. D. 2013. Arsenic-mediated activation of the Nrf2-Keap1 antioxidant pathway. J Biochem Mol Toxicol, 27, 99–105.

Love, M. I., Huber, W. & Anders, S. 2014. Moderated estimation of fold change and dispersion for RNA-seq data with DESeq2. Genome biology, 15, 1–21.

Lu, X., Xie, Q., Pan, X., Zhang, R., Zhang, X., Peng, G., Zhang, Y., Shen, S. & Tong, N. 2024. Type 2 diabetes mellitus in adults: pathogenesis, prevention and therapy. Signal Transduct Target Ther, 9, 262.

Mattaliano, M. D., Montana, E. S., Parisky, K. M., Littleton, J. T. & Griffith, L. C. 2007. The Drosophila ARC homolog regulates behavioral responses to starvation. Mol Cell Neurosci, 36, 211–21.

Mazumder, D. N. G., Ghosh, A., Majumdar, K. K., Ghosh, N., Saha, C. & Mazumder, R. N. G. 2010. Arsenic Contamination of Ground Water and its Health Impact on Population of District of Nadia, West Bengal, India. Indian Journal of Community Medicine, 35, 331–338.

Misra, J. R., Lam, G. & Thummel, C. S. 2013. Constitutive activation of the Nrf2/Keap1 pathway in insecticide-resistant strains of Drosophila. Insect Biochem Mol Biol, 43, 1116–24.

Moraru, A., Wiederstein, J., Pfaff, D., Fleming, T., Miller, A. K., Nawroth, P. & Teleman, A. A. 2018. Elevated Levels of the Reactive Metabolite Methylglyoxal Recapitulate Progression of Type 2 Diabetes. Cell Metab, 27, 926–934.e8.

Morimoto, R. I. 2011. The heat shock response: systems biology of proteotoxic stress in aging and disease. Cold Spring Harb Symp Quant Biol, 76, 91–9.

Mosher, J., Zhang, W., Blumhagen, R. Z., D’Alessandro, A., Nemkov, T., Hansen, K. C., Hesselberth, J. R. & Reis, T. 2015. Coordination between Drosophila Arc1 and a specific population of brain neurons regulates organismal fat. Dev Biol, 405, 280–90.

MUñIZ Ortiz, J. G., Opoka, R., Kane, D. & Cartwright, I. L. 2009. Investigating arsenic susceptibility from a genetic perspective in Drosophila reveals a key role for glutathione synthetase. Toxicological Sciences, 107, 416–426.

MUñIZ Ortiz, J. G., Shang, J., Catron, B., Landero, J., Caruso, J. A. & Cartwright, I. L. 2011. A transgenic Drosophila model for arsenic methylation suggests a metabolic rationale for differential dose-dependent toxicity endpoints. Toxicological Sciences, 121, 303–311.

Navas-Acien, A., Silbergeld, E. K., Pastor-Barriuso, R. & Guallar, E. 2008. Arsenic exposure and prevalence of type 2 diabetes in US adults. Jama, 300, 814–22.

Nemkov, T., Reisz, J. A., Gehrke, S., Hansen, K. C. & D’Alessandro, A. 2019. High-throughput metabolomics: isocratic and gradient mass spectrometry-based methods. High-Throughput Metabolomics: Methods and Protocols, 13–26.

ORGANIZATION, W. H. 2022. Guidelines for drinking-water quality: incorporating the first and second addenda, World Health Organization.

Öztürk-Çolak, A., Marygold, S. J., Antonazzo, G., Attrill, H., Goutte-Gattat, D., Jenkins, V. K., Matthews, B. B., Millburn, G., Dos Santos, G., Tabone, C. J., Perrimon, N., Gelbart, S. R., Broll, K., Crosby, M., Dos Santos, G., Falls, K., Gramates, L. S., Jenkins, V. K., Longden, I., Matthews, B. B., Seme, J., Tabone, C. J., Zhou, P., Zytkovicz, M., Brown, N., Antonazzo, G., Attrill, H., Goutte-Gattat, D., Larkin, A., Marygold, S., Mclachlan, A., Millburn, G., Pilgrim, C., Öztürk-Çolak, A., Kaufman, T., Calvi, B., Campbell, S., Goodman, J., Strelets, V., Thurmond, J., Cripps, R. & Lovato, T. 2024. FlyBase: updates to the <i>Drosophila</i> genes and genomes database. GENETICS, 227.

Pang, Z., Chong, J., Zhou, G., DE LIMA Morais, D. A., Chang, L., Barrette, M., Gauthier, C., Jacques, P.-É., Li, S. & Xia, J. 2021. MetaboAnalyst 5.0: narrowing the gap between raw spectra and functional insights. Nucleic acids research, 49, W388–W396.

Parker, G. H., Gillie, C. E., Miller, J. V., Badger, D. E. & Kreider, M. L. 2022. Human health risk assessment of arsenic, cadmium, lead, and mercury ingestion from baby foods. Toxicology reports, 9, 238–249.

Patro, R., Duggal, G., Love, M. I., Irizarry, R. A. & Kingsford, C. 2017. Salmon provides fast and bias-aware quantification of transcript expression. Nature methods, 14, 417–419.

Pederzolli, C. D., Mescka, C. P., Zandoná, B. R., De Moura Coelho, D., Sgaravatti, Â. M., Sgarbi, M. B., DE SOUZA Wyse, A. T., Duval Wannmacher, C. M., Wajner, M. & Vargas, C. R. 2010. Acute administration of 5-oxoproline induces oxidative damage to lipids and proteins and impairs antioxidant defenses in cerebral cortex and cerebellum of young rats. Metabolic brain disease, 25, 145–154.

Pederzolli, C. D., Sgaravatti, Â. M., Braum, C. A., Prestes, C. C., Zorzi, G. K., Sgarbi, M. B., Wyse, A. T., Wannmacher, C. M., Wajner, M. & Dutra-Filho, C. S. 2007. 5-Oxoproline reduces non-enzymatic antioxidant defenses in vitro in rat brain. Metabolic brain disease, 22, 51–65.

Podgorski, J. & Berg, M. 2020. Global threat of arsenic in groundwater. Science, 368, 845–850.

Puig, O. & Tjian, R. 2005. Transcriptional feedback control of insulin receptor by dFOXO/FOXO1. Genes Dev, 19, 2435–46.

Rahimi Kakavandi, N., Mousavi, T., Asadi, T., Moradi, A., Esmaeili, M., Habibian Sezavar, A., Nikfar, S. & Abdollahi, M. 2023. An updated systematic review and dose-response meta-analysis on the relation between exposure to arsenic and risk of type 2 diabetes. Toxicol Lett, 384, 115–127.

Ramos-Morales, P. & Rodriguez-Arnaiz, R. 1995. Genotoxicity of two arsenic compounds in germ cells and somatic cells of Drosophila melanogaster. Environmental and molecular mutagenesis, 25, 288–299.

Rand, M. D. 2010. Drosophotoxicology: the growing potential for Drosophila in neurotoxicology. Neurotoxicol Teratol, 32, 74–83.

Rand, M. D., Tennessen, J. M., Mackay, T. F. C. & Anholt, R. R. H. 2023. Perspectives on the <i>Drosophila melanogaster</i> Model for Advances in Toxicological Science. Current Protocols, 3.

Ratnaike, R. N. 2003. Acute and chronic arsenic toxicity. Postgraduate medical journal, 79, 391–396.

Rizki, M., Kossatz, E., VeláZquez, A., Creus, A., Farina, M., Fortaner, S., Sabbioni, E. & Marcos, R. 2006. Metabolism of arsenic in Drosophila melanogaster and the genotoxicity of dimethylarsinic acid in the Drosophila wing spot test. Environmental and molecular mutagenesis, 47, 162–168.

Rizki, M., Kossatz, E., Xamena, N., Creus, A. & Marcos, R. 2002. Influence of sodium arsenite on the genotoxicity of potassium dichromate and ethyl methanesulfonate: studies with the wing spot test in Drosophila. Environmental and molecular mutagenesis, 39, 49–54.

Rorsman, P. & Huising, M. O. 2018. The somatostatin-secreting pancreatic δ-cell in health and disease. Nat Rev Endocrinol, 14, 404–414.

Scanlan, J. L., Gledhill-Smith, R. S., Battlay, P. & Robin, C. 2020. Genomic and transcriptomic analyses in Drosophila suggest that the ecdysteroid kinase-like (EcKL) gene family encodes the ‘detoxification-by-phosphorylation’ enzymes of insects. Insect Biochemistry and Molecular Biology, 123, 103429.

Schalkwijk, C. G. & Stehouwer, C. D. A. 2020. Methylglyoxal, a Highly Reactive Dicarbonyl Compound, in Diabetes, Its Vascular Complications, and Other Age-Related Diseases. Physiol Rev, 100, 407–461.

Sehgal, A., Price, J. L., Man, B. & Young, M. W. 1994. Loss of circadian behavioral rhythms and per RNA oscillations in the Drosophila mutant timeless. Science, 263, 1603–6.

Sharma, V. K. & Sohn, M. 2009. Aquatic arsenic: toxicity, speciation, transformations, and remediation. Environment international, 35, 743–759.

Signes-Pastor, A. J., Carey, M. & Meharg, A. A. 2016. Inorganic arsenic in rice-based products for infants and young children. Food chemistry, 191, 128–134.

Singh, A. P., Goel, R. K. & Kaur, T. 2011. Mechanisms pertaining to arsenic toxicity. Toxicology international, 18, 87.

Singh, R., Singh, S., Parihar, P., Singh, V. P. & Prasad, S. M. 2015. Arsenic contamination, consequences and remediation techniques: a review. Ecotoxicology and environmental safety, 112, 247–270.

Soneson, C., Love, M. & Robinson, M. 2015. Differential analyses for RNA-seq: transcript-level estimates improve gene-level inferences [version 1; peer review: 2 approved]. F1000Research, 4.

Su, Y.-H., Mcgrath, S. P. & Zhao, F.-J. 2010. Rice is more efficient in arsenite uptake and translocation than wheat and barley. Plant and Soil, 328, 27–34.

Sykiotis, G. P. & Bohmann, D. 2008. Keap1/Nrf2 signaling regulates oxidative stress tolerance and lifespan in Drosophila. Developmental cell, 14, 76–85.

Taylor, V., Goodale, B., Raab, A., Schwerdtle, T., Reimer, K., Conklin, S., Karagas, M. R. & Francesconi, K. A. 2017. Human exposure to organic arsenic species from seafood. Science of the Total Environment, 580, 266–282.

Tchounwou, P. B., Yedjou, C. G., Udensi, U. K., Pacurari, M., Stevens, J. J., Patlolla, A. K., Noubissi, F. & Kumar, S. 2019. State of the science review of the health effects of inorganic arsenic: perspectives for future research. Environmental toxicology, 34, 188–202.

Thomas, D. J. 2009. Unraveling arsenic--glutathione connections. Toxicol Sci, 107, 309–11.

Upadhyay, M. K., Shukla, A., Yadav, P. & Srivastava, S. 2019. A review of arsenic in crops, vegetables, animals and food products. Food chemistry, 276, 608–618.

Van Der Pol, A., Gil, A., Silljé, H. H., Tromp, J., Ovchinnikova, E. S., Vreeswijk-Baudoin, I., Hoes, M., Domian, I. J., Van De Sluis, B. & Van Deursen, J. M. 2017. Accumulation of 5-oxoproline in myocardial dysfunction and the protective effects of OPLAH. Science translational medicine, 9, eaam8574.

Vincent, M. & Tanguay, R. M. 1982. Different intracellular distributions of heat-shock and arsenite-induced proteins in Drosophila Kc cells. Possible relation with the phosphorylation and translocation of a major cytoskeletal protein. J Mol Biol, 162, 365–78.

Vipat, S. & Moiseeva, T. N. 2024. The TIMELESS Roles in Genome Stability and Beyond. Journal of Molecular Biology, 436, 168206.

Vosshall, L. B., Price, J. L., Sehgal, A., Saez, L. & Young, M. W. 1994. Block in nuclear localization of period protein by a second clock mutation, timeless. Science, 263, 1606–9.

Wang, D., Kim, B. F., Nachman, K. E., Chiger, A. A., Herbstman, J., Loladze, I., Zhao, F.-J., Chen, C., Gao, A. & Zhu, Y. 2025. Impact of climate change on arsenic concentrations in paddy rice and the associated dietary health risks in Asia: an experimental and modelling study. The Lancet Planetary Health.

Wangler, M. F., Yamamoto, S., Chao, H.-T., Posey, J. E., Westerfield, M., Postlethwait, J., Network, M. O. T. U. D., Hieter, P., Boycott, K. M., Campeau, P. M. & Bellen, H. J. 2017. Model Organisms Facilitate Rare Disease Diagnosis and Therapeutic Research. Genetics, 207, 9–27.

Wu, C., Wilson, S., Walker, B., Dawid, I., Paisley, T., Zimarino, V. & Ueda, H. 1987. Purification and properties of Drosophila heat shock activator protein. Science, 238, 1247–53.

Yates, ANDREW D., Allen, J., Amode, R. M., Azov, A. G., Barba, M., Becerra, A., Bhai, J., CAMPBELL, Lahcen I., Carbajo Martinez, M., Chakiachvili, M., Chougule, K., Christensen, M., Contreras-Moreira, B., Cuzick, A., Da Rin Fioretto, L., Davis, P., De SILVA, Nishadi H., Diamantakis, S., Dyer, S., Elser, J., Filippi, C. V., Gall, A., Grigoriadis, D., Guijarro-Clarke, C., Gupta, P., Hammond-Kosack, KIM E., Howe, K. L., Jaiswal, P., Kaikala, V., Kumar, V., Kumari, S., Langridge, N., Le, T., Luypaert, M., Maslen, G. L., Maurel, T., Moore, B., Muffato, M., Mushtaq, A., Naamati, G., Naithani, S., Olson, A., Parker, A., Paulini, M., Pedro, H., Perry, E., Preece, J., Quinton-Tulloch, M., Rodgers, F., Rosello, M., Ruffier, M., Seager, J., Sitnik, V., Szpak, M., Tate, J., Tello-Ruiz, MARCELA K., Trevanion Stephen J., Urban, M., Ware, D., Wei, S., Williams, G., Winterbottom, A., Zarowiecki, M., Finn, ROBERT D. & Flicek, P. 2021. Ensembl Genomes 2022: an expanding genome resource for non-vertebrates. Nucleic Acids Research, 50, D996–D1003.

Yiwen, W., Xiaohan, T., Chunfeng, Z., Xiaoyu, Y., Yaodong, M. & Huanhuan, Q. 2022. Genetics of metallothioneins in Drosophila melanogaster. Chemosphere, 288, 132562.

Zdraljevic, S., Fox, B. W., Strand, C., Panda, O., Tenjo, F. J., Brady, S. C., Crombie, T. A., Doench, J. G., Schroeder, F. C. & Andersen, E. C. 2019. Natural variation in C. elegans arsenic toxicity is explained by differences in branched chain amino acid metabolism. eLife, 8, e40260.

Zhang, B., Egli, D., Georgiev, O. & Schaffner, W. 2001. The Drosophila homolog of mammalian zinc finger factor MTF-1 activates transcription in response to heavy metals. Mol Cell Biol, 21, 4505–14.

Zimarino, V., Wilson, S. & Wu, C. 1990. Antibody-mediated activation of Drosophila heat shock factor in vitro. Science, 249, 546–9.

