## Supplemental Figures for "Conserved Molecular Responses to Arsenite Exposure in *Drosophila melanogaster*"

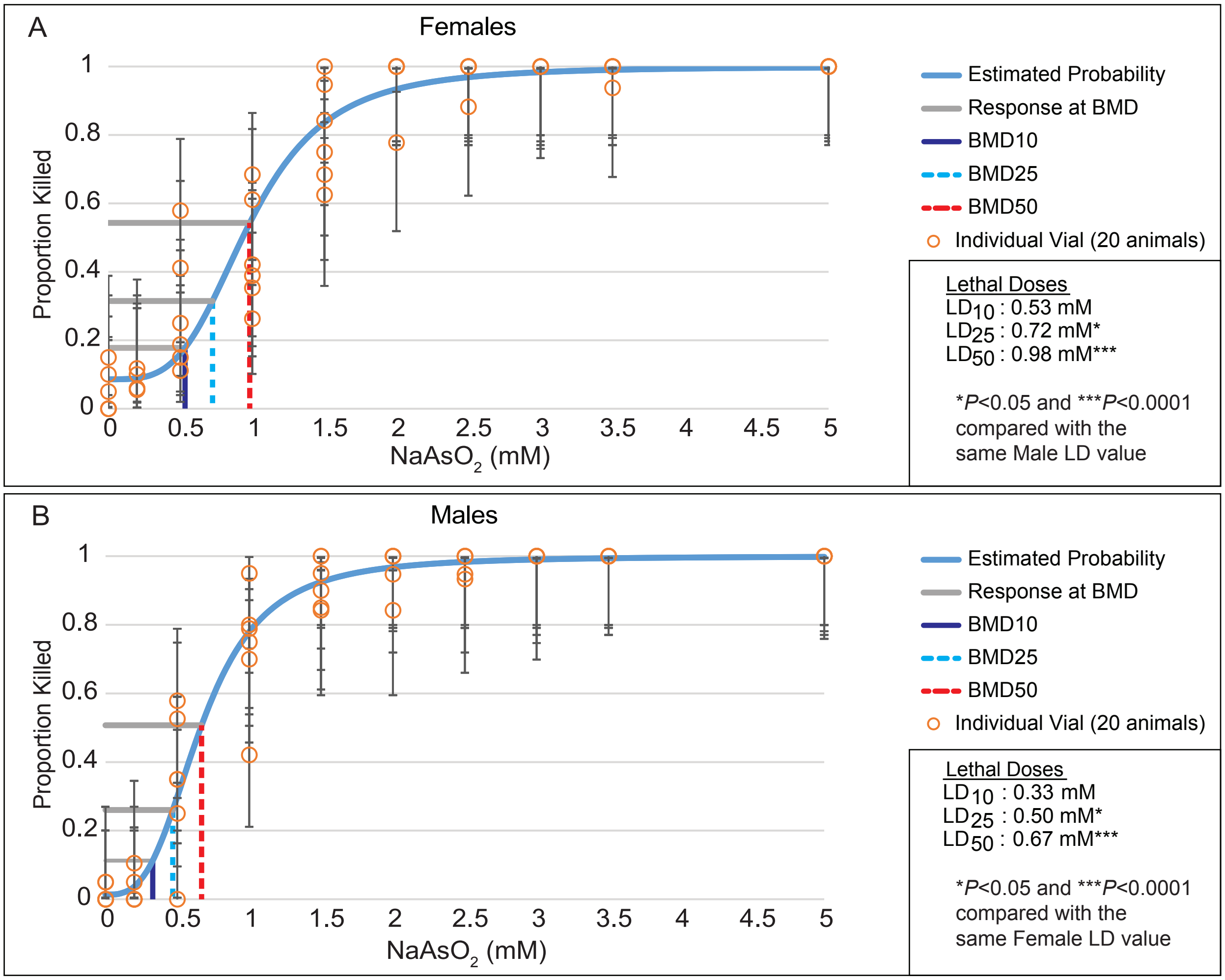


**Figure S1. Dose-response curves of *Drosophila* adult males and females exposed to $NaAsO_2$.** Dose-response curves illustrate the measured proportions of dead (A) female and (B) male Oregon-R flies at increasing concentrations of $NaAsO_2$ following a 48 hr exposure. Both the best fit line and the $LD_{10}$, $LD_{25}$, and $LD_{50}$ were calculated using a Dichotomous Hill Model. n=6 vials at each concentration with 20 flies per vial. (A,B – see text in lower right of panel) Male and female dose response curves were statistically analyzed with a global nonlinear regression model analysis using an extra sum-of-squares F test. This analysis revealed a significant difference in the $LD_{25}$ and $LD_{50}$ values between Oregon-R male and female flies ($LD_{10}$ F-statistic of $F(1,112) = 1.581$, $p = 0.2112$; $LD_{25}$ F-statistic of $F(1,112) = 5.586$, $p = 0.0198$; $LD_{50}$ F-statistic of $F(1,112) = 16.85$, $p < 0.0001$). However, the hillslope of the dose response curve was similar between males and females (F-statistic of $F(1,112) = 0.3659$, $p = 0.5465$).

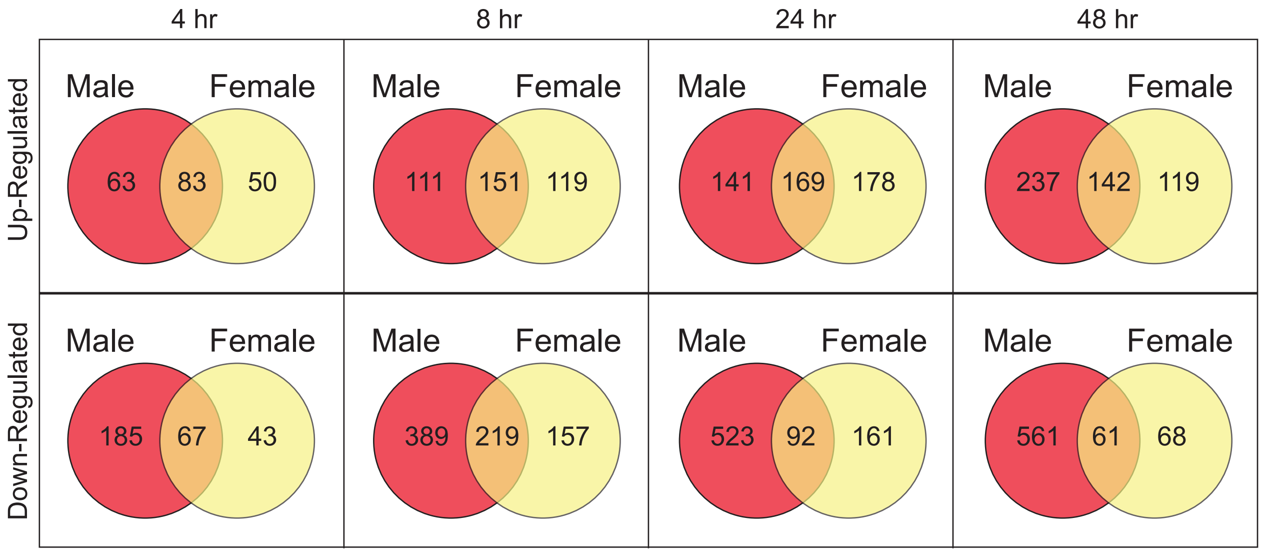


**Figure S2. A comparison of DEG in males and females exposed to 1.0 mM $NaAsO_2$.** Venn diagrams illustrating the overlap between up- and down-regulated DEGs in males and females at 4, 8, 24, and 48 hrs. Note that less than 50% of the DEGs at any given time point are shared between the two sexes. The 1 hr and 2 hr time points were excluded from this analysis due to the low number of DEGs.

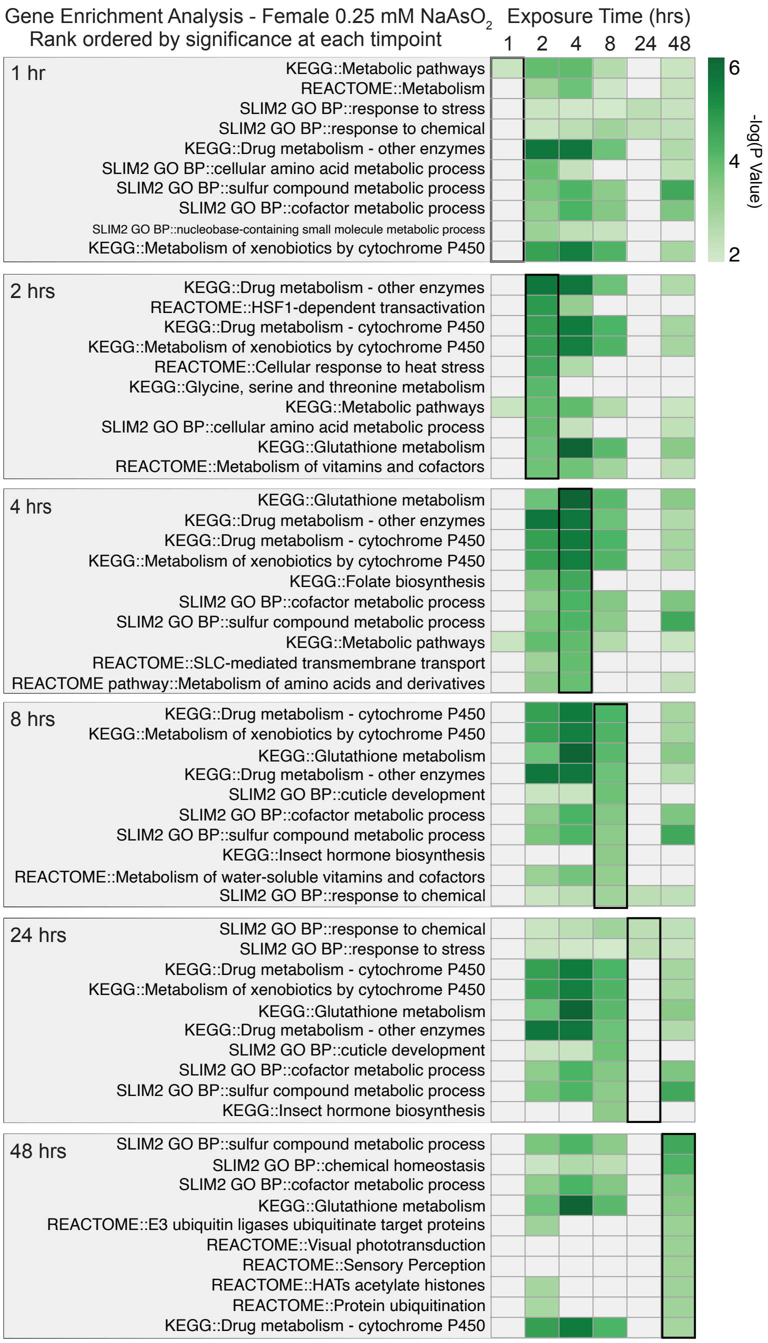


**Figure S3. Gene Set Enrichment Analysis of DEGs in female *Drosophila melanogaster* exposed to 0.25 mM $NaAsO_2$ across a 48-hour time course.** As in Figure 3, PANGEA was used to identify significantly overrepresented gene sets among differentially expressed genes (DEGs) at each exposure time point (see Methods and Table S3). DEGs from each time point were simultaneously analyzed for enrichment using four annotation sources: (i) SLIM2 GO Biological Process, (ii) FlyBase signaling pathways (experimental evidence), (iii) KEGG Pathway (*D. melanogaster*), and (iv) REACTOME Pathway (*D. melanogaster*). The top 10 enriched gene sets for each time point are presented with heatmaps of their rank-ordered adjusted *p*-value at the corresponding time (outlined heat map column), along with their significance (green gradient) or lack thereof (grey) at the other time points.

[illegible]

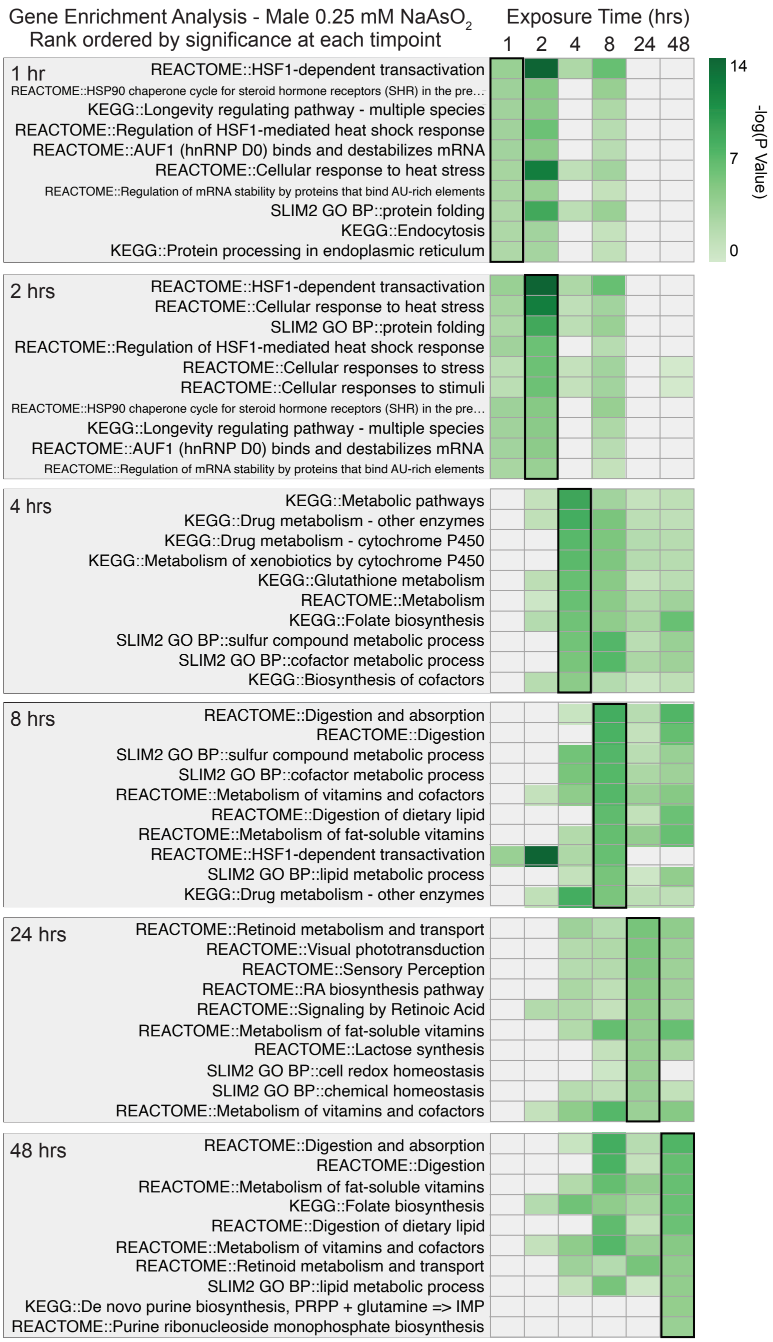


**Figure S4. Gene Set Enrichment Analysis of DEGs in male *Drosophila melanogaster* exposed to 0.25 mM $NaAsO_2$ across a 48-hour time course.** As in Figure 3, PANGEA was used to identify significantly overrepresented gene sets among differentially expressed genes (DEGs) at each exposure time point (see Methods and Table S5). The top 10 enriched gene sets for each time point are presented with heatmaps of their rank-ordered adjusted *p*-value at the corresponding time (outlined heat map column), along with their significance (green) or lack thereof (grey) at the other time points.

[illegible]

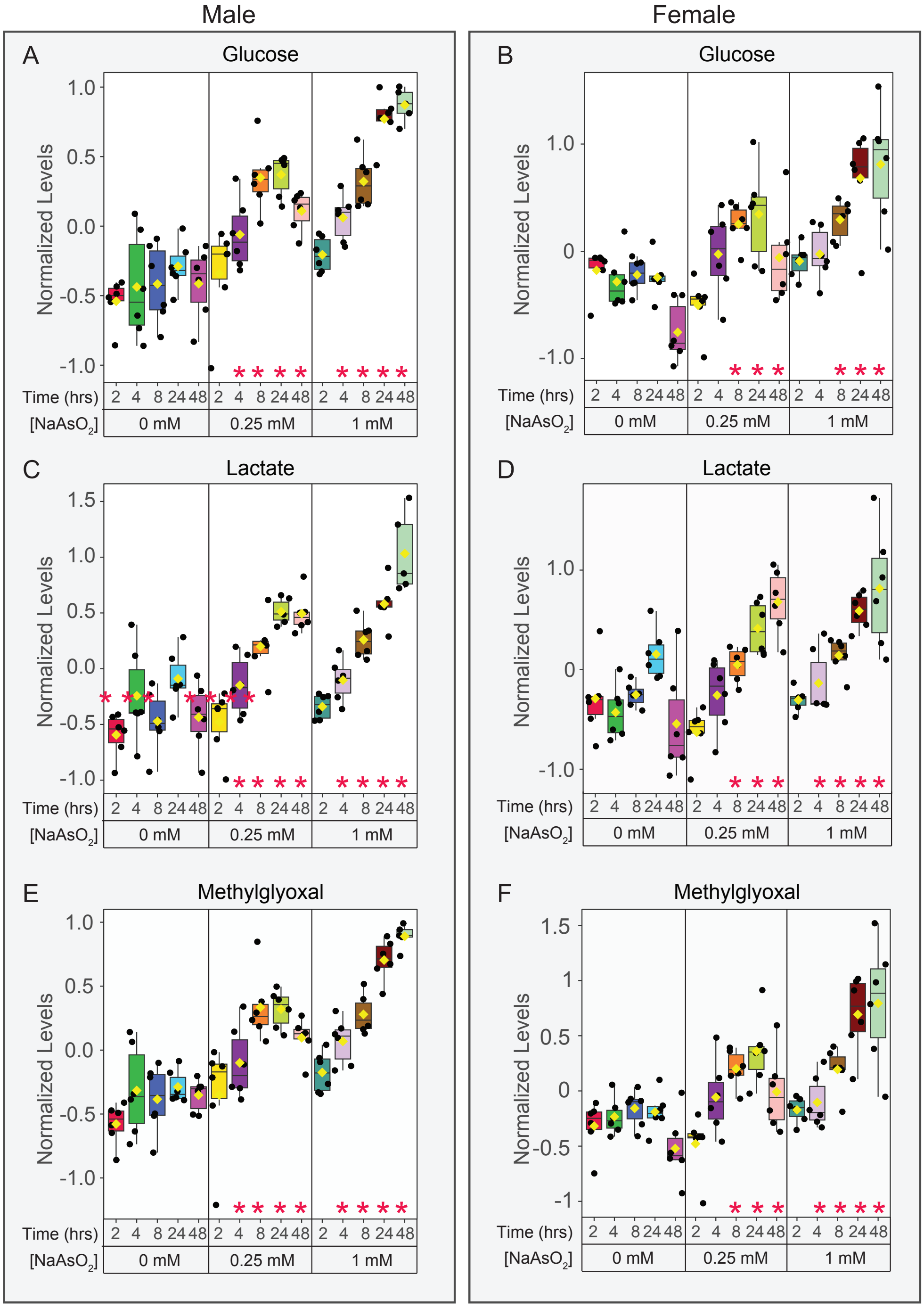


**Figure S5. $NaAsO_2$ exposure disrupts glycolytic metabolism in a dose- and time-dependent manner.** Box and whiskers plots displaying normalized levels of (A, B) glucose, (C, D) lactate, and (E, F) methylglyoxal in adult flies exposed to 0 mM (control), 0.25 mM, or 1.0 mM $NaAsO_2$ over a 48-hour time course. Analysis conducted with Metaboanalyst 6.0. Data were normalized to sample mass and processed with Log Transformation and Pareto scaling. Statistical analysis was conducted using ANOVA with FDR correction (FDR < 0.05), followed by Fisher's Least Significant Difference post hoc test (see Tables S20 and S21). **p* < 0.05 relative to untreated time-matched control. n=6 samples per condition.
